# State-dependent top-down and bottom-up processes in gamma-band (∼40 Hz) oscillations of the cat EEG

**DOI:** 10.64898/2026.07.20.739616

**Authors:** Santiago Castro-Zaballa, Joaquín González, Matías Cavelli, Pablo Torterolo

## Abstract

Cognitive processes rely on extensive thalamocortical and corticocortical recurrent interactions. The gamma frequency band (∼40 Hz) of the electroencephalogram (EEG) emerges from these interactions. Importantly, cognitive processing depends on the interplay between bottom-up sensory inputs and top-down influences from higher-order cortical areas. However, the extent to which gamma-band oscillations are associated with directional patterns of functional interactions remains unclear. Therefore, the aim of this study was to investigate the directionality of gamma-band information flow during wakefulness (W) and sleep, under spontaneous conditions and in response to auditory stimulation.

Cats were chronically implanted for polysomnographic recordings, with electrodes placed in multiple cortical and thalamic regions. Information-flow directionality was assessed using two complementary methods: (i) time-lag analysis of gamma-band amplitude envelopes between pairs of channels, and (ii) Granger Causality analysis of the same channel pairs.

During quiet W gamma-band oscillations exhibited a predominantly top-down directional organization, from higher- to lower-order cortical areas, as well as from cortical regions to thalamic nuclei. Following auditory stimulation, a prominent gamma response emerged between 0.5 and 1.5 s after stimulus onset. Within this time window, distinct patterns of gamma-band information flow were observed depending on the nature of the stimulus. Specifically, a bottom-up directional predominance was observed for simple auditory stimuli (clicks), whereas top-down processing prevailed for complex variable stimuli. In contrast, no consistent directionality of information flow was observed during either NREM or REM sleep, regardless of whether auditory stimulation was present.

These findings extend our understanding of gamma-band information-flow dynamics during wakefulness and sleep.

## 1. Introduction

Cognitive processes and consciousness depend on extensive, large-scale recurrent thalamocortical and corticocortical interactions (Redinbaugh *et al*., 2020; Schneider *et al*., 2020; Vinck *et al*., 2022). The gamma frequency band (30 to 100 Hz) of the electroencephalogram (EEG) arise as a result of these interactions and play a role in cognitive functions (Llinás & Ribary, 1993; Mashour, 2004). These oscillations have been implicated in the integration of spatially separated but temporally correlated neural events, leading to a cohesive perceptual experience (Cantero *et al*., 2004; Castro *et al*., 2013; Schneider *et al*., 2020).

Cortical and subcortical areas are interconnected through functional circuits, referred to as ’bottom-up’ and ’top-down’ based on the direction of the information they transmit. Bottom-up processing denotes the flow of sensory information ascending the cortical hierarchy, representing the physical attributes of the presented stimuli. This process may also trigger stimulus-driven attentional mechanism (Siegel *et al*., 2015; Siegle *et al*., 2021; Xiong *et al*., 2023). In other words, bottom-up processing is related to the flow of sensory information from the periphery to the thalamus, and then to the primary sensory cortex. Bottom-up information is processed in increasingly higher-order cortex, namely, in occipital, temporal, parietal association cortices, and finally in the prefrontal cortex.

Top-down is defined as the direction of information flow from higher-order to lower-order areas (Bastos, Litvak, *et al*., 2015; Bastos, Vezoli, Bosman, *et al*., 2015; Bastos *et al*., 2018, 2020; Vinck *et al*., 2022; Xiong *et al*., 2023). Top-down processing refers to the modulation of incoming sensory information by higher-order cognitive processes, including attention, goals, prior knowledge, and internal expectations. This type of information processing relies on internal knowledge or rules that have been acquired through learning and cannot be deduced solely based on the stimulus configuration presented to a subject at a given time (Xiong *et al*., 2023). Recurrent interactions between bottom-up and top-down streams may underlie flexible, context-dependent processing of sensory stimuli, as well as the formation of prediction and expectation; the interplay between these two processes is critical for different aspects of cognition (Vinck *et al*., 2022; Xiong *et al*., 2023).

Gamma oscillations during wakefulness (W) can occur spontaneously, or in response to sensory stimuli, such as visual (Murty & Ray, 2022; Pattisapu & Ray, 2023), auditory (Llinás & Ribary, 1993; Joliot, Ribary, & Llinas, 1994; Zanto *et al*., 2006; Ross, 2008; King *et al*., 2013) or olfactory (González *et al*., 2023). Sensory stimulation elicits two distinct types of gamma band activity: early or stimulus-coupled (evoked), and late or stimulus-uncoupled (induced) responses (Karakaş *et al*., 2001). Evoked responses are synchronized with the onset of the stimulus and can be analyzed by averaging the responses (Tallon-Baudry *et al*., 1996; Tallon-Baudry & Bertrand, 1999). These responses reflect the integration of the stimuli at the different levels of the sensory pathways and association cortices (Karakaş *et al*., 2001; Snyder & Large, 2004; Zanto *et al*., 2006; Ross, 2008). On the contrary, induced gamma activity is characterized by a loose temporal relationship with the onset of the stimulus. These signals are recorded with some fluctuation in their latency from one stimulus to the other, so they tend to cancel each other out in the average response (Tallon-Baudry *et al*., 1996; Tallon-Baudry & Bertrand, 1999). Induced gamma oscillations represent aspects less directly related to the stimuli, and more related to the cognitive aspects of sensory integration (Tallon-Baudry *et al*., 1996; Tallon-Baudry & Bertrand, 1999; Karakaş *et al*., 2001; Zanto *et al*., 2006). This signal is relevant to high-level cognitive processes, such as tasks that require the activation of visual or acoustic object representations (Tallon-Baudry *et al*., 1996; Tallon-Baudry & Bertrand, 1999; Karakaş *et al*., 2001). Karakaş et al. (2001) concludes that gamma oscillation is multifunctional; in the early time-window (evoked), gamma fulfills basically a sensory function, in the late window (induced) it fulfills perceptual-cognitive functions (Karakaş *et al*., 2001).

While evoked and induced response are better understood in the context of task related activity, less is known about how the direction of information is altered during different brain states. In mammals, three behavioral states can be distinguished: W, non-REM sleep (NREM) and rapid eye movement sleep (REM) (Mondino *et al*., 2022; Torterolo *et al*., 2022). Gamma activity is high during W and, although reduced, persists during sleep, particularly during REM sleep (Llinás & Ribary, 1993; Karakaş *et al*., 2006; Castro *et al*., 2013; Torterolo *et al*., 2019, 2022). Although there are studies in primates that have described the bottom-up and top-down directionality in the alpha/beta (8-30 Hz) and gamma (30-100 Hz) frequency bands activity during W (Bastos, Litvak, *et al*., 2015; Bastos, Vezoli, Bosman, *et al*., 2015; Siegel *et al*., 2015; Bastos *et al*., 2018, 2020; Xiong *et al*., 2023), the precise directionality of information flow encoded by gamma band oscillations during W remains unclear, and this process has not yet been explored during sleep.

In the present study, we investigated the directionality of low gamma-band (30-45 Hz) activity in the cat EEG, a frequency band that is particularly prominent in this species, using two complementary analytical approaches. One is straightforward; we quantified the direction of time shifts in the amplitude envelopes of the filtered low gamma oscillations of the EEG and thalamic electrogram channels. The second method was based on Granger Causality analysis. We observed that during quiet W (QW, without sensory stimulation), as well as in response to complex auditory stimuli (CVS, 200 ms random excerpts of a melody), top-down gamma activity prevails. In contrast, simple auditory stimuli (clicks) predominantly produces a bottom-up information flow. Interestingly, a predominant directionality is absent during both NREM and REM sleep.

## 2. Experimental procedures

### 2.1. Experimental animals

Six adult cats (3 males, 4.5 ± 1.5 kg) were included in this study. The animals were obtained from the Institutional Animal Care Facility. They were housed in individual enclosures under controlled light-dark cycles and temperature conditions, with *ad libitum* access to food. All experiments were approved by the Institutional Animal Care Commission (*Comisión Honoraria de Experimentación Animal de la Universidad de la República, and Comisión de Etica de Facultad de Medicina*), Protocol No: 070151-000013-22. Measures were taken to minimize pain, discomfort, or stress, and effort was made to employ the minimum number of animals necessary to generate reliable scientific data.

### 2.2. Surgical procedures

The animals were prepared for chronic polysomnographic recordings in semi-restricted conditions. Surgical and recordings procedures were those previously used by our group (Torterolo *et al*., 2016; Cavelli *et al*., 2020; Castro-Zaballa *et al*., 2024; Gallo *et al*., 2024). Briefly, the animals were implanted with electrodes in different neocortical areas under general anesthesia. A diagram showing the electrode positions is presented in Figure 1A. Electrodes were located in the ventral prefrontal cortex (Pfv, in the medial orbital gyrus), dorsolateral prefrontal cortex (Pfdl, in the lateral orbital gyrus), primary motor cortex (M1, in the posterior sigmoid gyrus), primary somatosensory cortex (S1, in coronal gyrus), associative or posterior parietal cortex (Pp, in the middle suprasylvian gyrus,), primary auditory cortex (A1, in middle ectosylvian gyrus), primary visual cortex (V1, in the posterior marginal gyrus). In addition, bipolar electrodes were placed in the visual thalamus (lateral geniculate nucleus, LGN) and, in two cats, in the auditory thalamus (medial geniculate nucleus, MGN). The anatomical placement of the electrodes followed the works of Stolzberg et al. (2017) and Pakozdy et al. (2015) and was guided by the stereotaxic atlas of Berman and Jones (1982) (Berman & Jones, 1982; Pakozdy *et al*., 2015; Stolzberg *et al*., 2017). The electrodes were soldered to connectors and fixed to the skull with acrylic cement. We also implanted a device that allows the head to be fixed to a stereotactic frame in a painless manner during the recording (see below).

**Figure 1.**
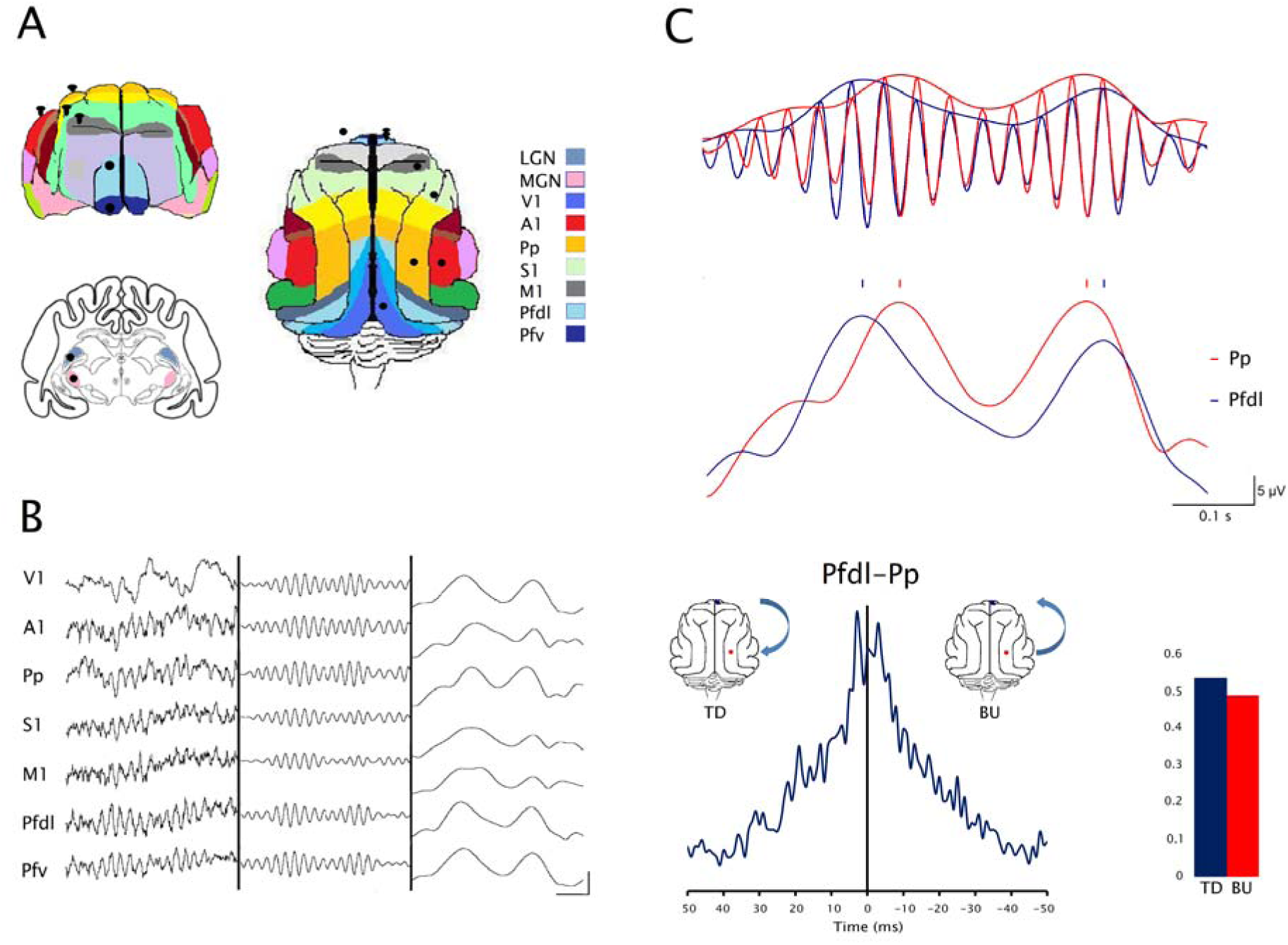
Method for analyzing the direction of the gamma information flow. A. Anterior and superior views of the cat brain, as well as a brain coronal section, showing the position of the recording electrodes over the cerebral cortex, the thalamus, and the reference electrode over the left frontal sinus. B. Raw recordings (left), band-pass filtered (30-45 Hz) recordings (center), and envelopes of gamma oscillations (right) from cortical areas recorded during quiet wakefulness. Calibration bars: 50 ms; 30 µV, 30 µV, and 10 µV, respectively. C. Quantification of the directionality of phase shifts of the envelopes of filtered gamma oscillations. Top. Filtered low gamma oscillations with its respective envelopes. Medium. Low gamma oscillations envelopes of Pfdl and Pp cortices with marks indicating the peak of the envelopes. Bottom left. Event-correlation histogram showing the temporal relationship between gamma peaks in the Pfdl and Pp cortices, computed from a 500-s recording window. Top-down (TD) and bottom-up (BU) events are labeled. Bottom right. Graph that shows the number of events (peaks of the envelopes of the gamma oscillations) in each directionality for the same time window. Pfv: Ventral prefrontal cortex; Pfdl: Dorsolateral prefrontal cortex; M1: Primary motor cortex; S1: Primary somatosensory cortex; Pp: Posterior parietal cortex; A1: Primary auditory cortex; V1: Primary visual cortex; MGN: medial geniculate nucleus; LGN: lateral geniculate nucleus

After the surgery, an analgesic was administered every 24 hours for three days. The incision margins were kept clean and topical antibiotics were applied daily. Once recovered from surgery, the animals were adapted to the experimental conditions for a period of no less than three weeks.

### 2.3. Experimental sessions

We recorded cortical and thalamic areas with monopolar electrodes (reference on the left frontal sinus) during the light phase. The electromyogram (EMG, with acutely placed bipolar electrodes on the skin over the dorsal neck muscles), electrocardiogram (precordial bipolar electrodes) and respiration (by means of a motion detector placed on the thorax and a thermistor in the nostrils) were also monitored.

As in our previous works, recordings were made under semi-restricted conditions (fixed head and body in a sleeping bag). To obtain a complete data set, cats are recorded daily during 4 hours, for 20 to 30 days. The bioelectrical signals were amplified (1000x), filtered (0.1-500 Hz), digitized (1024 Hz, 16 bits) and stored on a PC using Spike 2 software (Cambridge Electronic Design, CED). We performed auditory stimulation in 4 animals, through headphones placed in the external auditory canal of the animals. We used either clicks (1 ms bursts of white noise) or CVS, which were 200 ms excerpts randomly selected from a melody (Handel’s *Hallelujah*) using MATLAB, ensuring that each stimulus differed from the others. Clicks and CVS were applied at regular time intervals of 5 seconds. The intensity of the stimuli was approximately 60 dB SPL, and since the animals were highly habituated to the recording setup, usually the sounds did not awaken or prevent them from falling asleep.

The stimuli were delivered in six blocks of 120 events over 600 seconds (10 minutes) each, with 10-minute silent intervals between blocks; the total duration of the experiment was 2 hours. The auditory stimulation protocols were initiated remotely 30 minutes after the experimenter left the room. The stimuli were presented irrespective of the animals’ behavioral state.

### 2.4. Sleep and wakefulness

W, NREM and REM sleep were classified according to standard criteria (Ursin *et al*., 1981), and consistent with our previous studies. W was defined by the presence of an activated EEG and high-voltage nuchal EMG activity. NREM sleep by the presence of spindles and/or high-voltage low-frequency (0.5-3 Hz) waves in frontal and parietal cortex, as well as reduced tonic nuchal EMG activity. REM sleep by an activated EEG, atonia in the neck muscles, and the presence of ponto-geniculo-occipital (PGO) waves in the LGN recordings.

### 2.5. Selection of EEG time windows for analysis

We analyzed spontaneous or baseline gamma activity (without sensory stimulation) in six animals. This analysis was performed in continuous selected artifact free 500-second windows during each behavioral state (W, NREM and REM sleep). For each animal, we analyzed 144 windows in six recordings (24 windows/recording) per behavioral state.

We also evaluated stimulus-induced gamma activity during W and sleep for both clicks and CVS. For each behavioral state, gamma activity was analyzed in seven consecutive 500 ms-windows from stimulus onset up to 3.5 seconds post-stimulus. These analytical windows correspond to the early response (ER, 0-0.5 s; 1 window), the late gamma response (LGR, 0.5-1.5 s; 2 windows), and the inter-stimulus period (ISP, 1.5-3.5 s, 4 windows) (see Section 3.2 for the rationale of these time windows). The same time-windows from six recordings of four animals were concatenated before analysis. The results from these seven consecutive 500-ms analysis windows are typically presented after being collapsed into four broader windows: the ER, LGR, and two ISP windows (see for example Figure 4C).

### 2.6. Analysis of gamma power and coherence

Previous studies from our group characterized the power and coherence of spontaneous EEG activity in cats (Castro *et al*., 2013; Castro-Zaballa, Cavelli, González, Monti, *et al*., 2019; Castro-Zaballa, Cavelli, González, Nardi, *et al*., 2019). In the present study, we investigate stimulus-induced gamma activity during both W and sleep. We performed power spectrum analyses on EEG recordings with a temporal resolution of 0.25 seconds and a frequency resolution of 4 Hz, using the SleepScore script in Spike2 software. Analyses were conducted in the same seven analytical windows described above (from stimulus onset to 3.5 seconds post-stimulus) during both W and sleep. To minimize habituation effects, only the first 50 stimuli were included in the analysis. For each time window, power spectra were averaged and the mean baseline power of the corresponding behavioral state was subtracted to highlight increases in gamma power.

We ran coherence analysis between two EEG channels (derivations) with a temporal resolution of 0.5 seconds and a frequency resolution of 4 Hz, using a modified version of the COHER 1S script in Spike2 software. These analyses were also conducted for the first 50 stimuli during both W and sleep. For each behavioral state, each of the seven-time windows described above were concatenated, resulting in an analytical window of 25 seconds comprising fifty 0.5-second blocks (one per stimulus). We applied the Fisher z-transformation to normalize the coherence data. Subsequently, we averaged the z-transformed coherence values across all recordings and animals

### 2.7. Analysis of the directionality of information flow

In the time windows described in previous sections, we analyzed the directionality of gamma oscillations using two methods: the time lag of the gamma envelopes and Granger Causality analysis.

#### 2.7.1. Time lag of gamma amplitude envelopes

This method quantifies directionality based on shifts in the amplitude envelopes of filtered gamma oscillations (30-45 Hz), as illustrated in Figure 1B and C. First, gamma oscillations were isolated using a finite impulse response filter. Then, the root mean square (RMS) amplitude of the signal was calculated, and the peaks in the signal envelops were identified. Event correlation histograms of these peaks between two channels were generated with a temporal resolution of 1 millisecond and a time range of 50 milliseconds. Finally, for each time window, we counted the number of events corresponding to each phase direction, excluding those with phase shifts 0 (less than 1 millisecond). Finally, a “directionality index” was calculated as follow:

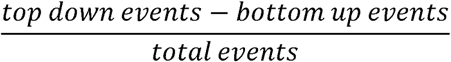

A positive value indicates a top-down predominance, while a negative value indicates a bottom-up predominance.

#### 2.7.2. Granger Causality

Granger Causality estimation is used to determine the magnitude and direction of the temporal relationship between simultaneously recorded signals (Granger, 1969; Seth, 2010; Hunt *et al*., 2019; Vinck *et al*., 2022; Xiong *et al*., 2023). It is a statistical hypothesis to determine whether a time series forecast or predict another. In the context of EEG analysis, Granger Causality between two simultaneously recorded channels allows to estimate the predominant direction of information flow between areas. The data was analyzed using MATLAB codes modified by the authors (Mathworks). We used a sampling rate of 1024 Hz and set the maximum model order for estimation to 20. Under these parameters, Granger Causality analysis evaluated whether one time series could predict another within a time window of approximately 20 ms (20 sampling points), which corresponds to the typical time scale of phase shifts in gamma oscillations.

### 2.8. Statistical analysis

Prior to statistical analyses, data normality was evaluated using the Shapiro-Wilk test, and homogeneity of variances was assessed using Levene’s test. Thereafter, power, coherence, Granger Causality and the directionality index values obtained during W, NREM and REM sleep were compared by one-way ANOVA with Bonferroni *post hoc* test. The same test was used to compare these parameters obtained in the different time-windows post stimuli. We also performed Student t tests to compare the number of bottom-up Vs. top-down time shifts gamma envelopes for each state. In all the cases, the result was considered significant with p < 0.05/n of comparisons (Bonferroni correction).

## 3. Results

### 3.1. Neocortical gamma top-down direction predominates during quiet wakefulness

Under QW (without sensory stimulation), Granger Causality and the analysis of the gamma oscillations envelopes indicate that the main direction of information flow in the gamma band was top-down. Figure 2A illustrates gamma oscillation directionality in an exemplary dataset from associative cortical areas, demonstrating a predominant top-down information flow from Pfdl to Pp. Figure 2A and B also show that the predominance of the top-down flow of information was absent during both NREM and REM sleep, and significantly different than QW.

**Figure 2.**
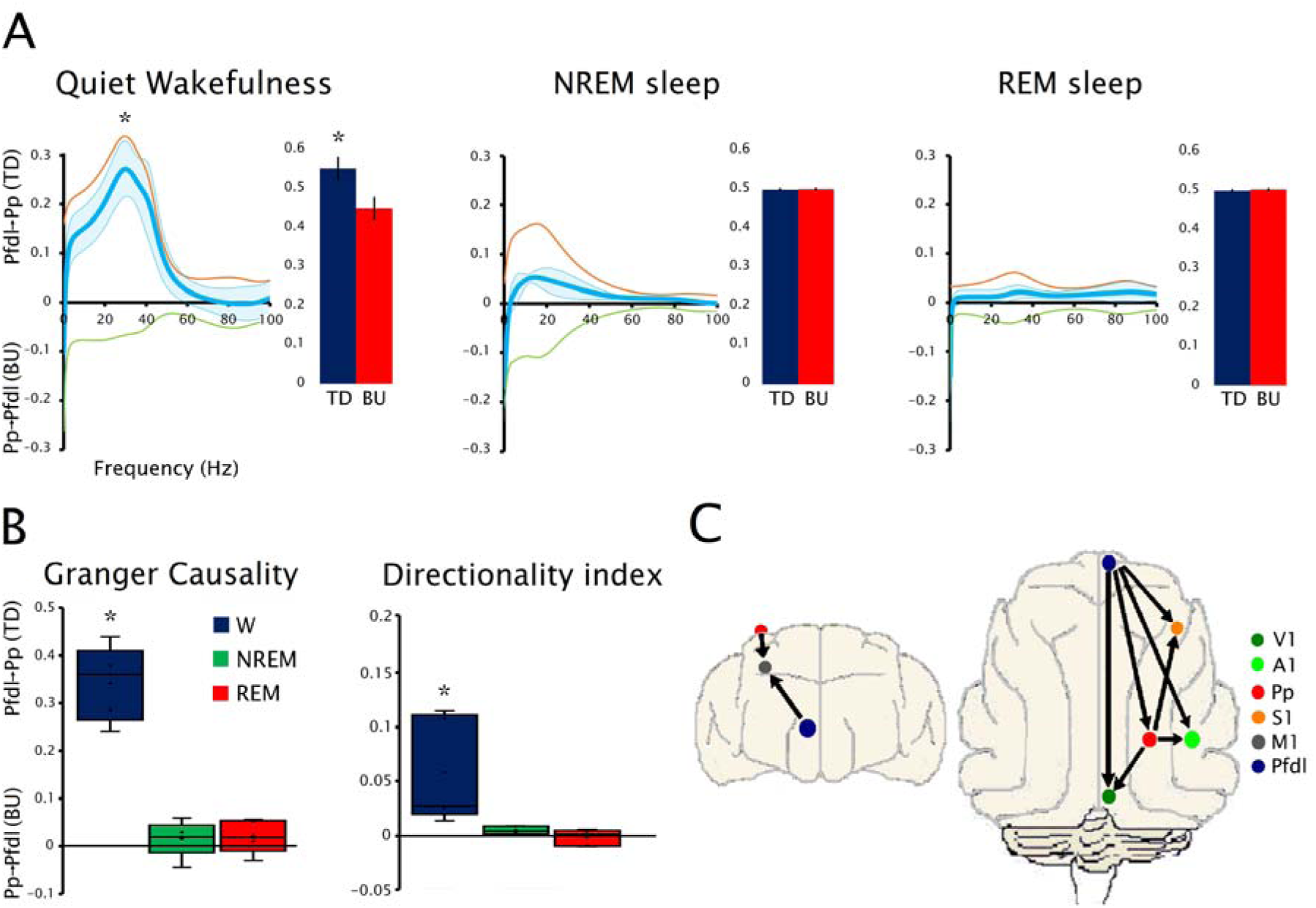
**Gamma directionality during basal conditions (without sensory stimulation)**. A. Mean and standard error of Granger Causality spectrum analysis during quiet wakefulness, NREM and REM sleep. The orange line shows the top-down, while the green line the bottom-up directionality. The light blue line illustrates the difference (subtraction) between both directionalities. These analyses are accompanied by graphs that show the number of events (peaks of the envelopes of the gamma oscillations) in each direction. B. Box-plot exhibiting the results of Granger Causality and directionality index during sleep for Pfdl and Pp derivation. C. Schematic diagram of the main sense of the gamma information flow. The colored circles indicate the position of the surface electrodes on the cerebral cortex, while the arrows show the significant predominant directionality of the gamma frequency band during wakefulness. Only the significant directions of the information flow are shown. All the analyses were conducted in six cats. Asterisks (*) show significance (ANOVA and Bonferroni *post hoc*) with p < 0.05. Pfv: Ventral prefrontal cortex; Pfdl: Dorsolateral prefrontal cortex; M1: Primary motor cortex; S1: Primary somatosensory cortex; Pp: Posterior parietal cortex; A1: Primary auditory cortex; V1: Primary visual cortex. TD: top-down; BU: bottom-up.

Figure 2C synthesizes the significant directionality during QW. The main direction of information flow in the gamma band was from Pfdl to Pp and from Pfdl to primary cortices (S1, M1, A1, and V1). Additionally, a predominant directionality was observed from Pp to the primary cortices. The analyses for all derivations during QW and sleep are shown in the Supplementary Figure 1. Across all channel combinations, the directional predominance observed in gamma oscillations during W was lost during sleep.

### 3.2. Gamma response to auditory stimulation

To investigate the direction of gamma-band information flow ("bottom-up" vs. "top-down") triggered by auditory stimuli, we employed two sound protocols: clicks and CVS. First, we characterized the gamma activity induced by these stimuli during W. As shown in Figure 3A, both types of stimuli applied during W elicit gamma oscillations in different cortical areas that are prominent between 0.5 and 1.5 seconds after stimulation. We refer to the gamma oscillations in this temporal window as the "late gamma response" (LGR).

**Figure 3.**
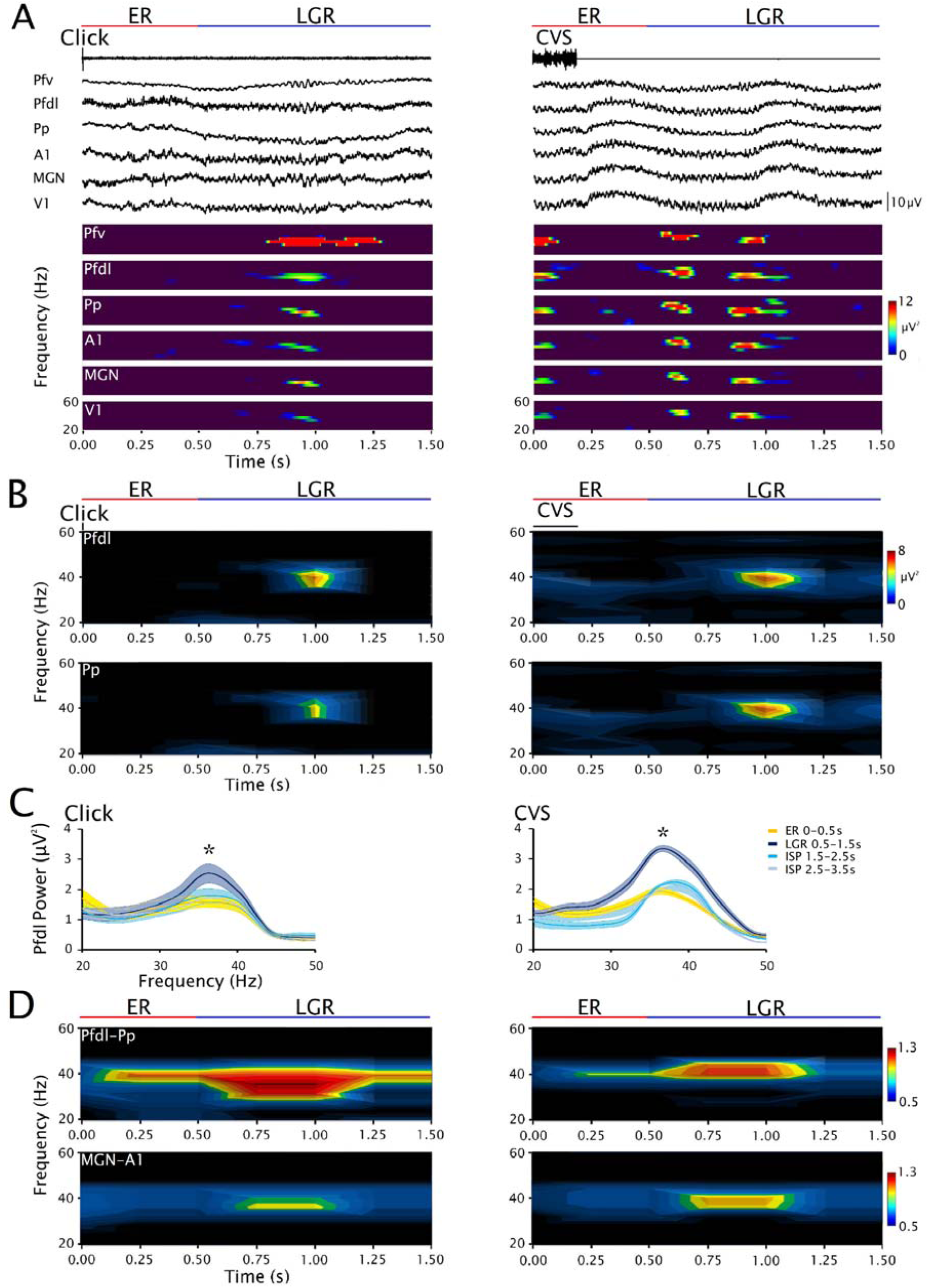
Gamma response induced by auditory stimulation. A. Raw recordings and spectrograms following clicks and complex variable stimuli (CVS), showing the late gamma response (LGR), from the following areas of the right hemisphere: ventral prefrontal cortex (Pfv), dorsolateral prefrontal cortex (Pfdl), posterior parietal cortex (Pp), primary auditory cortex (A1), primary visual cortex (V1) and medial geniculate nucleus (MGN). B. Average of the spectrograms from the Pfdl and Pp following clicks (1200 events) or CVS (1200 events) stimulation. These analyses were conducted during wakefulness in 4 animals. C. Mean and standard error of average gamma power of the Pfdl cortex in the four-time windows following click or CVS stimuli. Asterisks (*) show significance (ANOVA and Bonferroni *post hoc*) in comparison to the other time windows (p < 0.05). D. Average of Pfdl-Pp and MGN-A1 Źcoherograms following clicks or CVS stimulation.

Since the LGR diminishes (habituates) with repeated stimulation, we analyzed only the first 50 stimuli of each stimulation series. We analyzed six recordings per animal (4 animals). Thus, the analyses during W were based on EEG responses to a total of 1200 click stimuli and 1200 CVS presentations. Figure 3B presents the averaged spectrograms of the stimulus-induced responses, in two representative cortical areas, the Pfdl and Pp, where the LGR is evident.

Figure 3C presents the average power of the Pfdl cortex in four post-stimulus time windows: 0-0.5 seconds, corresponding to the ER associated with the evoked potentials (see Supplementary Figure 2); 0.5-1.5 seconds, corresponding to the LGR and two subsequent windows (1.5-2.5 and 2.5-3.5 s) following the LGR. These last two windows correspond to the ISP and can be considered as control. During LGR induced by either clicks or CVS, gamma power was higher than during ER and the ISP. On the contrary, gamma power during ER and the ISP were similar. The gamma power increment during the LGR was significant in all cortical and thalamic areas recorded (data not shown).

Gamma band coherence during LGR induced by both clicks and CVS was also higher than during ER and ISP in all cortical and thalamo-cortical derivations. Figure 3D shows examples of Z’coherence for Pfdl-Pp and MGN-A1 derivations. Since gamma power and coherence increased only during the LGR, the directionality analysis was performed only for this time window (see below).

We also analyzed the gamma activity following auditory stimuli during sleep. A total of 1200 events were analyzed during NREM (both clicks and CVS). However, only 138 click events and 126 CVS events were analyzed during REM sleep, as this behavioral state was shorter in duration than the others; nevertheless, at least one REM sleep episode was analyzed for each animal. Interestingly, neither click nor CVS stimulation altered gamma power or coherence during the time window corresponding to the LGR in either NREM or REM sleep, across all analyzed channels. An example of the Pfdl power during the LGR following clicks and CVS stimulation during W and sleep are shown in Supplementary Figure 3.

### 3.3. Neocortical gamma bottom-up direction predominates in the processing of clicks

LGR directionality following click stimulation was assessed during W. Figure 4A reveals a predominant bottom-up process from Pp to Pfdl, utilizing both measurements, Granger Causality and the gamma envelope event correlation. This bottom-up direction was absent when the signals were analyzed either before (during the ER) or in the ISP as shown in the temporal analysis of the Granger Causality for the Pfdl and Pp derivation of Figure 4B and C. Note in Figure 4B that the bottom-up direction extends to lower frequency bands. Figure 4D summarizes the significant directionality following click stimulation during W. A significant predominance was observed only from Pp to Pfdl and from A1 to Pfdl (the results for all derivations are shown in Supplementary Figure 4).

**Figure 4.**
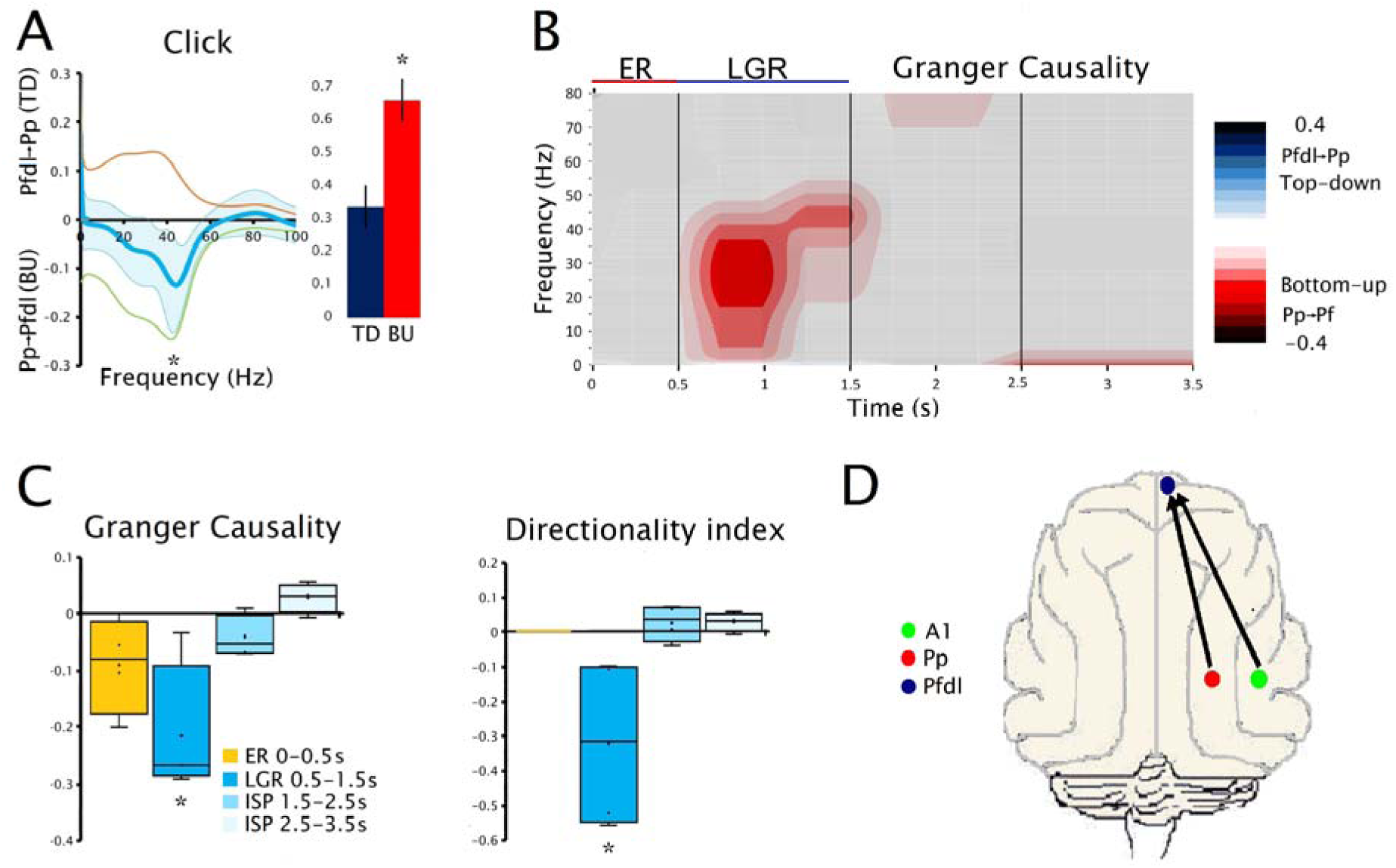
Directionality in late gamma response (LGR) induced by click stimuli. A. Left. Mean and standard error of Granger Causality analysis of the Pfdl and Pp derivation during the LGR (0.5 to 1.5 seconds after the stimuli) following click stimuli. Right. Graph that shows the number of events peaks (envelopes of the gamma oscillations) in each direction. B. Temporal analysis of the Granger Causality of the Pfdl and Pp cortices following click stimuli. C. Box-plot of Granger Causality and directionality index at different time windows following the stimulus. D. Schematic diagram of the main direction of the gamma information flow. The colored circles indicate the position of the surface electrodes on the cerebral cortex and arrows show the significant predominant directionality in the gamma frequency band during W. Asterisks (*) show significance (ANOVA and Bonferroni *post hoc*) compared to the other time windows (p < 0.05). TD: top-down; BU: bottom-up; Pfdl: Dorsolateral prefrontal cortex; A1: Primary auditory cortex; V1: Primary visual cortex. All the analyses were conducted in four cats.

LGR directionality following click stimulation was also analyzed during NREM and REM sleep (Supplementary Figure 4). No predominance in the direction of gamma oscillations was observed.

### 3.4. Neocortical gamma top-down direction predominates in the processing of CVS

Figure 5A shows the directionality of the CVS-induced LGR in the Pfdl-Pp derivation, revealing a clear predominance of top-down processing. This top-down directionality was only present in the LGR, but not in the ER or in the interstimulus period as it is observable in the Granger Causality temporal analysis of Figure 5B and in the box-plots of Figure 5C.

**Figure 5.**
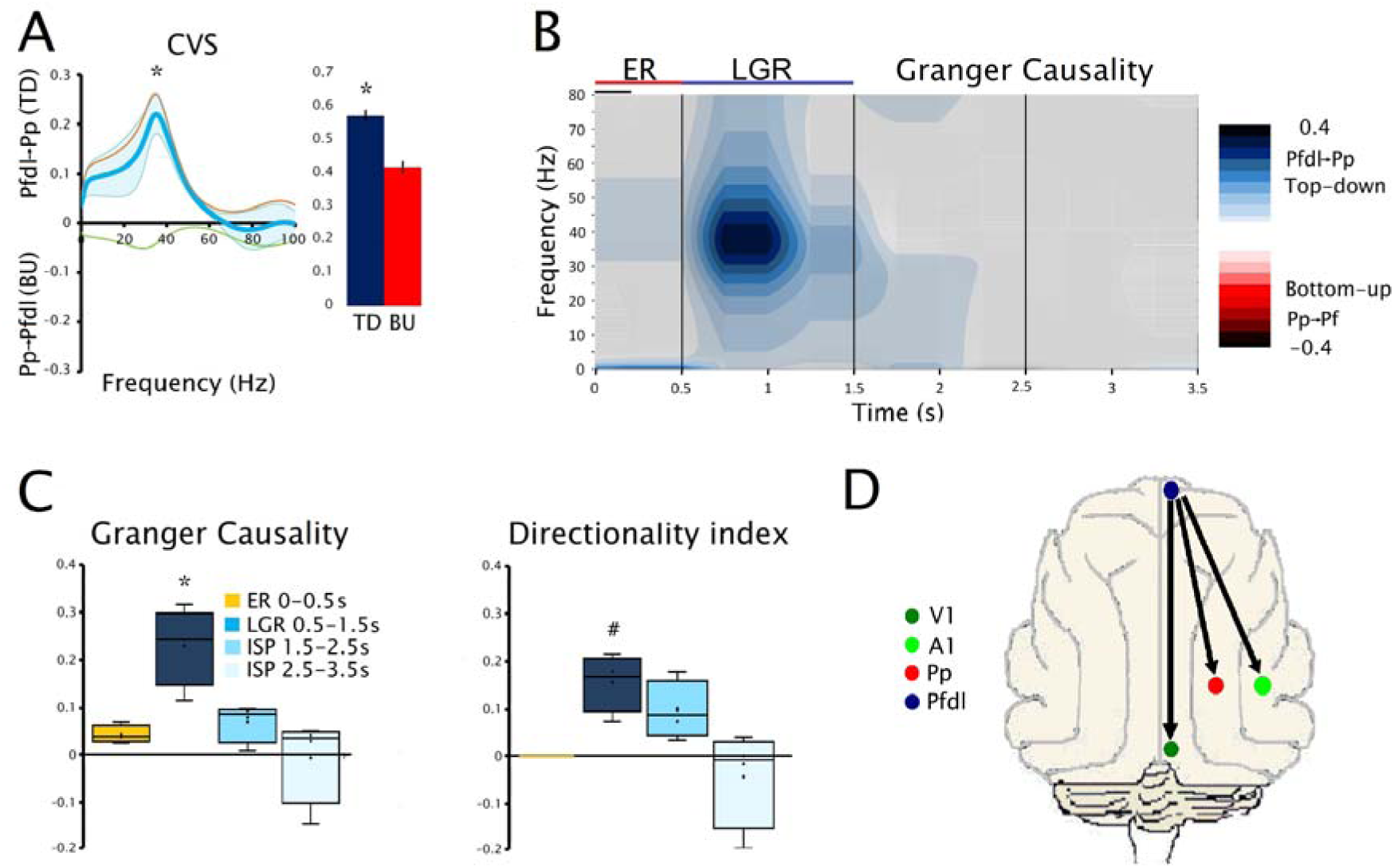
Directionality in late gamma response (LGR) induced by CVS stimuli. A. Left. Mean and standard error of Granger Causality analysis of the Pfdl and Pp cortices during the LGR (0.5 to 1.5 seconds after the stimuli) following CVS stimuli. Right. Graph that shows the number of events peaks (envelopes of the gamma oscillations) in each direction. B. Temporal analysis of the Granger Causality of the Pfdl and Pp derivation following CVS stimuli. C. Box-plot of Granger Causality and directionality index at different time windows following the onset of the stimulus. D. Schematic diagram of the main direction of the gamma information flow. The colored circles indicate the position of the surface electrodes on the cerebral cortex and arrows show the significant predominant directionality in the gamma frequency band during W. Asterisks (*) show significance (ANOVA and Bonferroni post hoc) with all the other time windows (p < 0.05) and numeral (#) shows significance Vs. ER and the second ISP time window. TD: top-down; BU: bottom-up; Pfdl: Dorsolateral prefrontal cortex; A1: Primary auditory cortex; Pp; posterior parietal cortex; V1: Primary visual cortex. All the analyses were conducted in four cats.

Figure 5D summarizes the significant directionality of the LGR following CVS stimulation during W. There was a predominant top-down directionality, originating from Pfdl and projecting to Pp, A1, and V1 (the results for all the derivations are shown in Supplementary Figure 5). Notably, these top-down directionalities were not observed during sleep (Supplementary Figure 5).

### 3.5. Gamma directionality between the thalamus and cortex

We also explored the gamma flow direction between the thalamus and the neocortex. First, we explored the directionality between the MGM and A1. Figure 6A shows that during QW and following clicks or CVS stimulation, the predominant direction is top-down; i.e., from A1 to MGN. Neither bottom-up nor top-down directionalities were observed between MGN and neocortical areas other than A1 (not shown).

**Figure 6.**
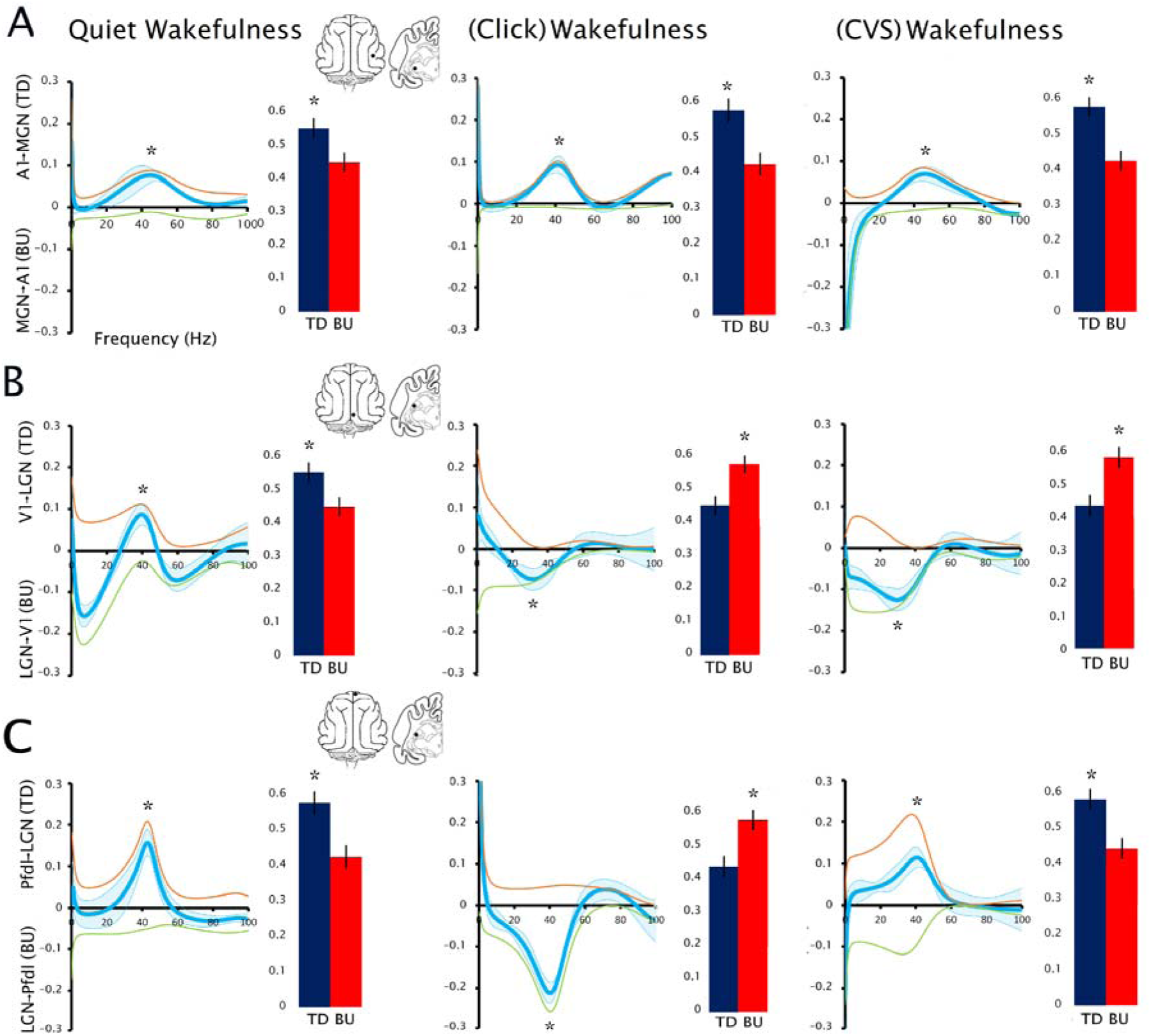
Directionality between neocortex and thalamus. A. Mean and standard error of Granger Causality spectrum of the A1 and MGN derivation during quiet wakefulness (without sensory stimulation), and during the late gamma response (0.5 to 1.5 seconds after the stimuli) induced by click or CVS stimuli. These analyses are accompanied by graphs that show the number of events of the peaks of the envelopes of the gamma oscillations in each direction. B. The same analysis for the LGN-V1 derivation. C. The same analysis for the PFdl-V1. Insets: Anterior and superior views of the cat brain showing the position of the analyzed recording electrodes over the cerebral cortex and thalamus. Asterisks (*) show significance (ANOVA and Bonferroni *post hoc*) with all the other time windows (p < 0.05). Pfdl: Dorsolateral prefrontal cortex; M1: Primary motor cortex; A1: Primary auditory cortex; V1: Primary visual cortex; MGN: medial geniculate nucleus; LGN: lateral geniculate nucleus. TD: top-down; BU: bottom-up. All the analyses were conducted in two cats.

We also explored the gamma flow direction between the visual thalamus (LGN) and cortical areas. We found that during QW there was a predominant top-down directionality; i.e., from V1 to LGN (Figure 6B), and from Pfdl to LGN (Figure 6C). Regarding the LGR directionality following sound stimuli during W, we found a predominant bottom-up directionality (from LGN to V1) both for clicks and CVS (Figure 6B). In contrast, while a bottom-up directionality was observed from the LGN to Pfdl following click stimulation, a top-down predominance was found for CVS stimulation (from Pfdl to LGN) (Figure 6C).

The gamma directionalities between thalamic nuclei and cortical areas are lost during both NREM and REM sleep (Supplementary Figure 6).

No significant directionalities were found during either QW, following auditory stimulation, or during sleep, between the LGN and other cortical areas, or between both thalamic nuclei (data not shown).

## 4. Discussion

The aim of our study was to investigate the dynamics of the information flow within the gamma band of the EEG during both W and sleep. We conducted this study using two analytical approaches: Granger Causality and gamma envelope correlation. We considered the directionality to be either bottom-up or top-down when the results obtained from both methods were significant. The results were consistent across methods; whenever a predominant directionality was observed, it was captured by both analytical approaches.

Our results show that during QW (without sensory stimulation), the predominant direction of the gamma band information flow among cortical areas or between cortical areas and thalamic nuclei were top-down.

We also analyzed the cortical directionality of gamma oscillations in response to auditory stimulation, focusing on what we termed "late gamma response", where gamma power and coherence was most prominent. The direction of the neocortical gamma oscillations during the LGR depended on the type of sound stimuli. When a simple, repetitive sound such as click, was applied, the bottom-up direction prevailed. In contrast, following the stimulation with complex and variable stimuli such as CVS, the top-down direction dominated.

In addition, we analyze the interactions between the thalamus and the cortex following sound stimulation during W. We found that the LGR induced by both types of stimuli is characterized by a predominant top-down directionality in the A1-MGN derivation. On the contrary, there was a prevailing bottom-up directionality between the LGN and V1 for both types of sounds. In the LGN-Pfdl derivative, we found a predominant bottom-up directionality in response to click stimulation, whereas a top-down type of processing predominated following CVS.

Finally, during NREM and REM sleep, cortico-cortical and thalamo-cortical bottom-up and top-down processes were balanced, both under basal conditions (without sensory stimulation) and following clicks or CVS stimuli.

### 4.1. Technical considerations

Recordings were performed in cats adapted to head-restrained conditions. One advantage is that the EEG is not affected by posture or movement, and the presence of artifacts is reduced, which is an important factor given that Granger Causality analysis is highly sensitive to artifacts.

In addition, this model exhibits strong gamma oscillations (30-45 Hz) in the EEG during W, with minimal contamination from extracranial noise (Castro *et al*., 2013). While electromyogenic artifacts is a common challenge for gamma band analysis in humans (Hipp & Siegel, 2013), we have previously demonstrated that spontaneous gamma oscillations recorded in cats during cataplexy, a waking state characterized by complete muscle atonia and, therefore, free of muscle artifacts, have the same characteristics as those observed during natural W (Torterolo *et al*., 2016). Hence, we ruled out the possibility that gamma oscillations were produced by muscle electrical activity. Another argument supporting the notion that we are recording genuine gamma oscillations is the variable latency observed between gamma wave peaks across the different cortices recorded simultaneously; this latency was proportional to the distance between the recording electrodes (Castro *et al*., 2013). In fact, the time shift of low gamma oscillations between different cortical pairs correlated with the distance between the recording electrodes (Supplementary Figure 7). This finding would not be expected if the recorded potentials originated from an extracerebral source, and also argues against volume conduction as a major contributor to our findings. Moreover, although saccadic eye movements in humans can generate gamma oscillations in frontal derivations (Yuval-Greenberg *et al*., 2008; Hipp & Siegel, 2013), the lack of coherence between the EEG and the electrooculogram, together with the absence of gamma coherence among cortical areas during REM sleep, when saccadic eye movements are most frequent, indicates that these movements did not contribute to the generation of gamma activity in our model (Castro *et al*., 2013).

### 4.2. Gamma oscillations in the feline EEG during wakefulness

During W, gamma oscillations around 40 Hz are widely distributed across the cat’s neocortex. These oscillations are organized in bursts lasting 200-500 ms, display a spindle-like morphology, and show high coherence across cortical areas (Castro *et al*., 2013). This gamma oscillations (which are also called beta by some authors) are very pronounced in the frontoparietal cortex when a cat is immobile, alert, and displaying behavior suggesting focused attention toward a target in its environment (Bouyer *et al*., 1981, 1983; Montaron *et al*., 1982; Rougeul-Buser, 1994). In addition, the gamma band increases its power and coherence when a novel stimuli is introduced to the cat (Bouyer *et al*., 1981; Castro *et al*., 2013; Castro-Zaballa, Cavelli, González, Nardi, *et al*., 2019; Castro-Zaballa *et al*., 2024).

It is important to note that gamma band activity should not be confused with gamma oscillations (Ichim *et al*., 2024). The former refers to any signal with spectral power in the gamma frequency range; however, this does not necessarily indicate the presence of a true oscillation or rhythmic process (Yuval-Greenberg *et al*., 2008; Buzsáki & Wang, 2012; Castro *et al*., 2013; Ardelean *et al*., 2023; Ichim *et al*., 2024). In the present study, we focus on true gamma oscillations confined to the 30-45 Hz range within the gamma band, as demonstrated in the cat by Gallo et al. (2024), using FOOOF (Fitting Oscillations and One-Over F) technique, that allows to parameterize power spectra and extract their periodic and aperiodic components (Gallo *et al*., 2024).

As mentioned in the Introduction, gamma oscillations can be induced by sensory stimulation; however, this type of induced gamma oscillation has not been previously studied in cats. Therefore, we characterized those using well-established approaches: computing power spectrograms for each individual trial, averaging power across trials, and performing coherence analysis (Tallon-Baudry & Bertrand, 1999; Bertrand *et al*., 2001).

We found that that auditory stimulation elicited an increase in gamma activity that began approximately 0.5 seconds after stimulus onset and persisted up to 1.5 seconds (Figure 3). As expected for an induced-response, and in contrast to the transient or steady-state evoked responses (Karmos *et al*., 2002), the gamma response was characterized by a loose temporal relationship with the stimulus (Bertrand *et al*., 2001; Karakaş *et al*., 2001). Another distinctive feature of the induced activity is the synchronization of the signal between distant electrodes, suggesting that widely separated cortical areas can become synchronized (Tallon-Baudry *et al*., 1996; Rodriguez *et al*., 1999; Bertrand *et al*., 2001). Notably, the LGR were coherent not only across different cortical areas, but also with the thalamus (Figure 3D). Following Karakaş et al. (2001), we termed this signal as the late gamma response (LGR).

Since the LGR represents the only period with a significant increase in gamma power and coherence, we focused our directionality analysis on this time window. Note that a signal similar to what we termed LGR has been described in scalp recordings from humans during a simple auditory detection task (Jokeit & Makeig, 1994). According to these authors, the 40 Hz signal increased between 200 and 400 ms after stimulus onset with respect to the pre-stimulus level. With frequent standard tones, complex stimuli and rare deviant tones, the induced gamma response peaks occurred at 500-800 msec, and remains up to 1000 ms or more (Bertrand *et al*., 2001).

### 4.3. Gamma top-down and bottom-up interaction during wakefulness

Although gamma oscillations are generated locally in each cortical and thalamic area (Buzsáki & Wang, 2012); our interpretation is that top-down directionality correspond to gamma oscillations of a higher-order area influencing the generation of gamma oscillations in lower-order areas, and bottom up directionality correspond to gamma oscillations of a lower-order area influencing the generation of gamma oscillations in higher-order areas. These influences give rise to delayed gamma-band synchronization between cortical areas (Bastos, Vezoli, & Fries, 2015; Fries, 2015; Ichim *et al*., 2024).

Previous studies in macaques have reported delayed gamma-band synchronization between visual areas V1 and V4 (Vinck *et al*., 2010, 2013; Maris *et al*., 2013), as well as a bidirectional directionality (analyzed by Granger Causality) for these frequency band between these areas (Bosman *et al*., 2012; Bastos, Vezoli, Bosman, *et al*., 2015). The role of top-down influences on gamma-band activity has also been explored by selectively suppressing feedback projections from the higher-order visual area 21a to V1 (Ye *et al*., 2026). However, the dynamics of spontaneous low gamma oscillations and those induced by different types of auditory stimuli have not yet been studied, either in cats or in other species.

Hence, we first explored the direction of the gamma-information flow during QW, i.e., without sensory stimulation. We found that there is a predominant top-down directionality, from higher order associative cortical areas to primary cortical areas, as well as from Pfdl to Pp (Figure 2).

We then explored the directionality of the LGR. The LGR induced by click stimuli has a predominant bottom-up directionality, from A1 to Pfdl as well as from Pp to Pfdl. Interestingly, the difference between bottom-up and top-down events was high. For example, the directionality index for Pp to Pfdl interactions show more than 75% of the events being bottom-up (Figure 4) and from A1 to Pfdl, up to 80% of the events being bottom-up (Supplementary Figure 4).

In contrast, LGR induced by CVS has a predominant top-down directionality, from Pfdl to Pp, A1, and V1 cortex. One interesting issue is the fact that there was a significant top-down directionality from Pfdl cortical area to V1 area, despite the auditory modality of stimuli (Figure 5D, Supplementary Figure 5). Even if it cannot be determined from the present experimental design, this result suggests that the animals direct their attention to both auditory and visual cues. Since each CVS differs from the previous one, the animals likely seek visual information to identify the source and nature of the novel sounds. In this regard, it has been reported that the prefrontal cortex and parietal cortex specifically encodes cross-modal representations, which emerges during tasks requiring the association of information across sensory modalities (Zhao & Ku, 2018; Gu *et al*., 2020; Park *et al*., 2025).

Regarding other sensory modalities, previous studies in macaques and humans have used Granger Causality to analyze the ER (associated with the evoked potential) in cortical visual areas during visual attention tasks. They show that 20-35 Hz oscillations, which the authors call high beta, was stronger in the top-down direction, and high gamma activity (80 Hz) was stronger in the bottom-up direction (Bastos, Litvak, *et al*., 2015; Bastos, Vezoli, & Fries, 2015; Michalareas *et al*., 2016; Bastos *et al*., 2018, 2020; Vinck *et al*., 2022; Qin *et al*., 2023). As commented before, our results showed that low gamma power did not increase during the ER; therefore, we focused on the LGR. In spite of this, we observed that Granger Causality scores during the ER were an order of magnitude smaller than those during the LGR (data not shown). Note that evoked potentials are broadband signals that can include gamma activity; however, they may not reflect true gamma oscillations. In contrast, the LGR during W consists of narrow-band oscillations around 40 Hz, sharing the same characteristics as the spontaneous gamma rhythm (Bouyer *et al*., 1981, 1983; Joliot, Ribary, & Llinás, 1994; Rougeul-Buser, 1994; Tallon-Baudry *et al*., 1996; Tallon-Baudry & Bertrand, 1999; Castro *et al*., 2013; Gallo *et al*., 2024). Hence, the interpretation of the gamma in these two time-windows should be different. Finally, it is noteworthy that the LGR was observed only during the first 50-80 stimuli and disappeared with habituation; animals frequently fell asleep thereafter despite continued stimulation. This pattern further suggests that the LGR is associated with a cognitive process rather than with simple sensory activation.

### 4.4. Functional interactions between the thalamus and the cortex

Low gamma oscillations have been recorded in the thalamus in the cat. These oscillations occurred and vanished simultaneously at thalamic and cortical sites with a high coherence between them (Bouyer *et al*., 1981; Rougeul-Buser, 1994). Here we extend the study of the gamma oscillations exploring the directionality between the thalamus and the cortex.

Regarding the auditory thalamus, the MGN receives auditory information from the inferior colliculus and have strong bilateral connection with A1 (Winer, 1984; Smith *et al*., 2019). An interesting anatomical feature is that feedback projections are larger than their feedforward counterparts (Pontes *et al*., 1975; Kimura *et al*., 2005). This feedback projections from A1 to MGN are believed to serve distinct gating functions in auditory attention (Kimura *et al*., 2005; Mellott *et al*., 2014). We found a predominant top-down directionality from A1 to MGN in the gamma oscillations during QW, as well as in the LGR induced by either clicks or CVS (Figure 6A).

It is important to highlight that there was not found a significant directionality between neither Pfdl and MGN, nor between Pp and MGN, which suggest that Pfdl cortex is not directly involved in top-down control of MGN as it does with A1.

Regarding the visual thalamus, we found a predominant top-down directionality from V1 to LGN during QW and a predominant bottom-up directionality from LGN to V1 in the LGR in response to click and CVS stimuli. We also found a strong predominance of top-down directionality between Pfdl cortex and NGL in both QW and in the LGR to CVS, while there was a predominant bottom-up directionality in the LGR to click. This suggest that Pfdl exercises a direct top-down control on the NGL when the animal is looking for visual cues to understand auditory stimuli that is interested in.

The difference in processing in the auditory and visual thalamus may be explained by variations in their respective connectivity. The prefrontal cortex (Pf) does not have neither direct input nor output projections to MGN. This cortex receives projections from the MGN through A1, and project towards the MGN through the thalamic reticular nucleus via the basal forebrain; however, the extent of these connections are unclear (Hockley & Malmierca, 2024). On the other hand, the Pf has stronger indirect connections to LGN via the mediodorsal thalamus (Griffiths *et al*., 2022), which might explain the predominance of top-down directionality between Pfdl cortex and LGN in both QW and in response to CVS.

Additionally, it has been suggested that the reduction in V1 activity observed during multisensory stimulation may originate from bottom-up signals from the LGN (Freeman *et al*., 2002; Baier *et al*., 2006), probably by activating inhibitory interneurons in V1. This aligns with the predominant bottom-up directionality from LGN to V1 observed following both clicks and CVS, but not during QW.

### 4.5. Top-down and bottom-up processing during wakefulness and sleep

Gamma oscillations has been associated with cognitive processing and consciousness during W (Torterolo *et al*., 2019; Ichim *et al*., 2024). In this regard, our results revealed that gamma oscillation have a predominant top-down flow from higher-order associative to primary cortical areas, with a strong directional predominance from Pfdl to Pp. These results support the concept that the frontoparietal network is important for the cognitive functions during W (Siegel *et al*., 2015; Sormaz *et al*., 2018; ElShafei *et al*., 2019).

Like in QW, the LGR induced by CVS showed a predominant top-down directionality, with a strong bias from Pfdl to Pp. Hence, processing complex and engaging stimuli involves functional interactions between associative prefrontal and parietal cortices (Siegel *et al*., 2015; Sormaz *et al*., 2018; ElShafei *et al*., 2019). We also found a robust top-down directionality from the Pfdl to A1. In fact, the prefrontal cortex exerts control over auditory processing through top-down modulation of the auditory pathway, influencing both perception and behavioral responses to sounds. This modulation has been implicated in processes such as deviance detection, attention, avoidance, and fear conditioning (Hockley & Malmierca, 2024; Hockley *et al*., 2025). In addition, dorsomedial and ventromedial prefrontal cortex are involved in modulation by top-down attention of gamma activity related to distracting sounds (ElShafei *et al*., 2019). Interestingly, whereas bottom-up processing predominated during the LGR to clicks, top-down processing predominated during the response to CVS. This finding suggests that CVS may engage the animals’ attention to a greater extent than simple click stimulation.

Profound changes in cognitive function occur during the transition from W to sleep. Conscious awareness is lost during deep NREM sleep, whereas REM sleep is characterized by vivid dreaming and represents a distinctive cognitive state that shares several features with psychotic experiences. (Hobson & Friston, 2012; Torterolo *et al*., 2019; Mutti *et al*., 2024; Tononi *et al*., 2024). Gamma oscillations are also modified during sleep. Gamma power and coherence decreases during NREM sleep in the cat, while gamma coherence is almost absent during REM sleep although gamma power has QW values (Castro *et al*., 2013; Cavelli *et al*., 2017; Castro-Zaballa, Cavelli, González, Nardi, *et al*., 2019; Castro-Zaballa *et al*., 2024). Hence, we expected that gamma directionally would be altered during sleep. In fact, we demonstrate that during both NREM and REM sleep in either basal condition, or following click or CVS stimulation, the predominance of top-down or bottom-up in gamma directionality disappear. This result was observed for both cortico-cortical interaction as well as for the interaction between the thalamus and the cortex.

In the thalamus, an oscillatory mode is associated with NREM sleep, that is characterized by a decreased ability to transfer incoming sensory inputs and by long-lasting periods of inhibition interrupted by burst discharges (Llinás & Paré, 1991). Auditory stimuli during NREM sleep shows evoked potentials characterized by a deep negative deflection lasting about half-a-second, which correspond to a K-complex or slow oscillations that represents an isolated cortical down-state, electrical silence or off-period. This cortical down-state that follows auditory stimuli may prevent the occurrence of induced gamma oscillations (Cash *et al*., 2009; Bellesi *et al*., 2014; Laurino *et al*., 2019; González *et al*., 2021; Cavelli *et al*., 2022).

Like W, thalamocortical cells show a relay mode during REM sleep. In this mode, an increased synaptic responsiveness, tonic discharge and short periods of inhibition are present in this neurons (Llinás & Paré, 1991). During REM sleep auditory stimulation produces an early, time-locked response (approximately 150 ms post stimulus) that can include gamma activity, but this sensory-perceptual processing is not followed by a clear LGR (Llinás & Ribary, 1993; Karakaş *et al*., 2006). Even though evoked potential studies indicate that the thalamocortical system is responsive to sensory input during REM sleep, gamma activity was neither reset by sensory stimuli nor enhanced upon stimulation during this state (Llinás & Ribary, 1993). Interestingly, sensory inputs of the external world are largely prevented from reaching conscious awareness during this state (Llinás & Paré, 1991). The lack of a clear gamma-band directionality during REM sleep may be related to this reduced processing of external sensory information.

### 4.6. Conclusions

In the present report, we found that during QW, the directional patterns of functional interactions in the cortical gamma band is primarily top-down. Following auditory stimulation, top-down flow between cortical areas dominates the LGR to complex stimuli, while bottom-up flow prevails for simple stimuli. Additionally, the direction from the cortex to the thalamus during the LGR differs according to the stimulus and the analyzed areas. Interestingly, the directional predominance disappears during both NREM and REM sleep. Thus, this report extend our understanding of the organization of the interactions of gamma oscillations in cognitive functions during W and sleep.

## Acknowledgements

This study was financed by CSIC I+D Groups Program 2022, CSIC I+D 2024 and ANII FCE 180456. All authors were also supported by the Uruguayan Sistema Nacional de Investigadores (SNI).

## CRediT (Contributor Roles Taxonomy) author statement

**Santiago Castro-Zaballa**: Conceptualization, Methodology, Validation, Formal analysis, Investigation, Data Curation, Writing- Original draft preparation, Visualization, Project administration, funding acquisition.

**Joaquín González**: Software, Validation, Writing - Review & Editing, Visualization.

**Matías Cavelli**: Methodology, Software, Validation, Writing - Review & Editing.

**Pablo Torterolo**: Conceptualization, Methodology, Validation, Writing - Review & Editing, Visualization, Supervision, Project administration, funding acquisition.

## Ethical statement

All experiments were approved by the Institutional Animal Care Commission (*Comisión Honoraria de Experimentación Animal de la Universidad de la República, and Comisión de Etica de Facultad de Medicina*), Protocol No: 070151-000013-22. Measures were taken to minimize pain, discomfort, or stress, and effort was made to employ the minimum number of animals necessary to generate reliable scientific data. All animal experiments were conducted in accordance with procedures authorized by the Bioethics Committee.

## Data availability statement

The data that support the findings of this study are available from the corresponding author upon reasonable request.

## Abbreviations

A1: primary auditory cortex
BU: bottom-up
CVS: complex variable sound
EEG: electroencephalogram
ER: early response
ISP: inter-stimulus period
LGR: late gamma response
LGN: lateral geniculate nucleus
M1: primary motor cortex
MGN: medial geniculate nucleus
NREM: non-REM sleep
Pfdl: dorsolateral prefrontal cortex
Pfv: ventral prefrontal cortex
Pp: posterior parietal cortex
QW: quiet wakefulness
REM: rapid eye movements sleep
RMS: root mean square
S1: primary somatosensory cortex
TD: top-down
V1: primary visual cortex
W: wakefulness

