## Supplementary Figures for "State-dependent top-down and bottom-up processes in gamma-band (∼40 Hz) oscillations of the cat EEG"

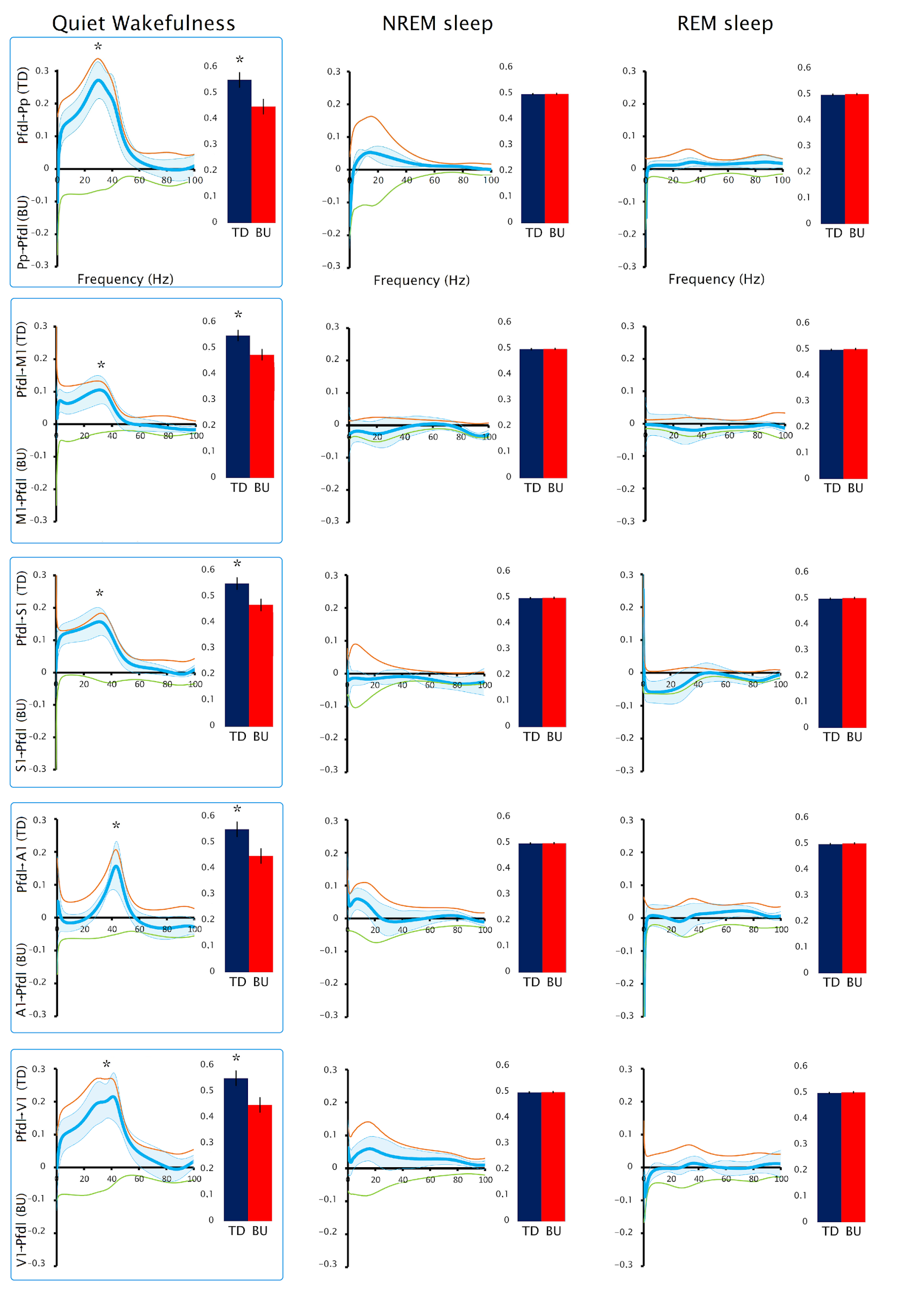

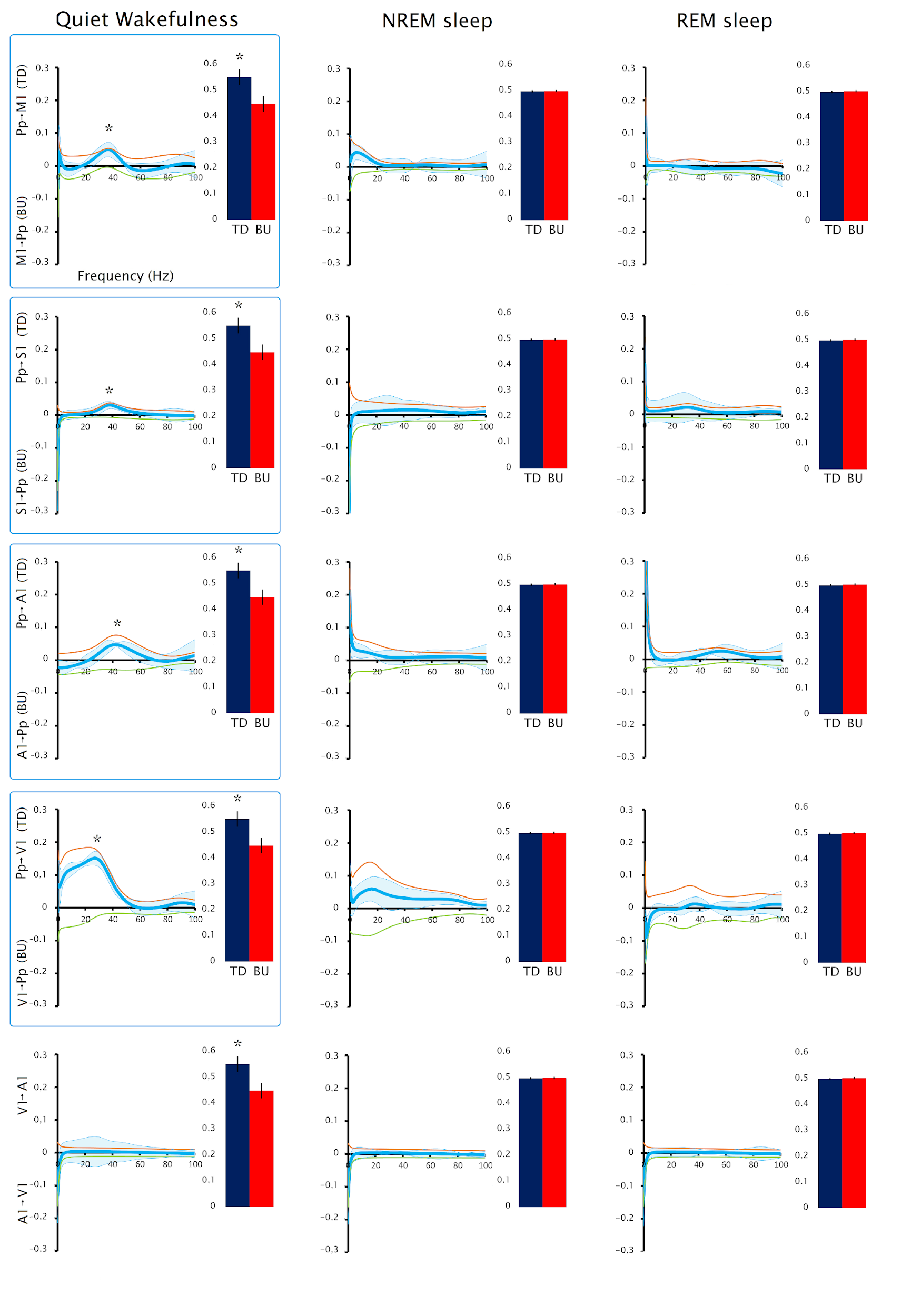

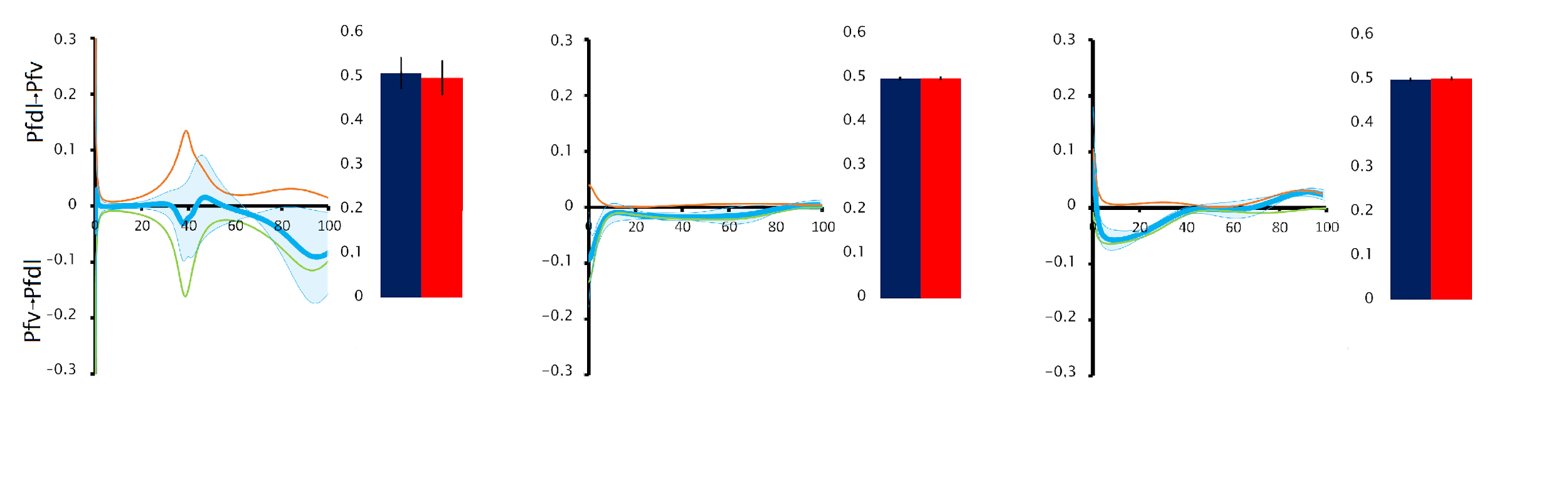

**Supplementary Figure 1.** **Gamma directionality during wakefulness and sleep for all the cortical derivatives during baseline conditions (without sensory stimulation).** Mean and standard error of Granger Causality spectrum analysis during wakefulness and sleep for all the cortical combinations during baseline conditions. The orange line shows the top-down, while the green line the bottom-up directionality. The light blue line illustrates the difference (subtraction) between both directionalities. These analyses are accompanied by graphs that show the number of events of the peaks of the envelopes of the gamma oscillations in each direction. All the analyses were conducted in six cats. Asterisk (*) shows significance (ANOVA and Bonferroni *post hoc*) with p < 0.05. EEG derivations showing statistically significant differences are outlined by boxes. Pfv: Ventral prefrontal cortex; Pfdl: Dorsolateral prefrontal cortex; M1: Primary motor cortex; S1: Primary somatosensory cortex; Pp: Posterior parietal cortex; A1: Primary auditory cortex; V1: Primary visual cortex. TD: top-down; BU bottom-up.

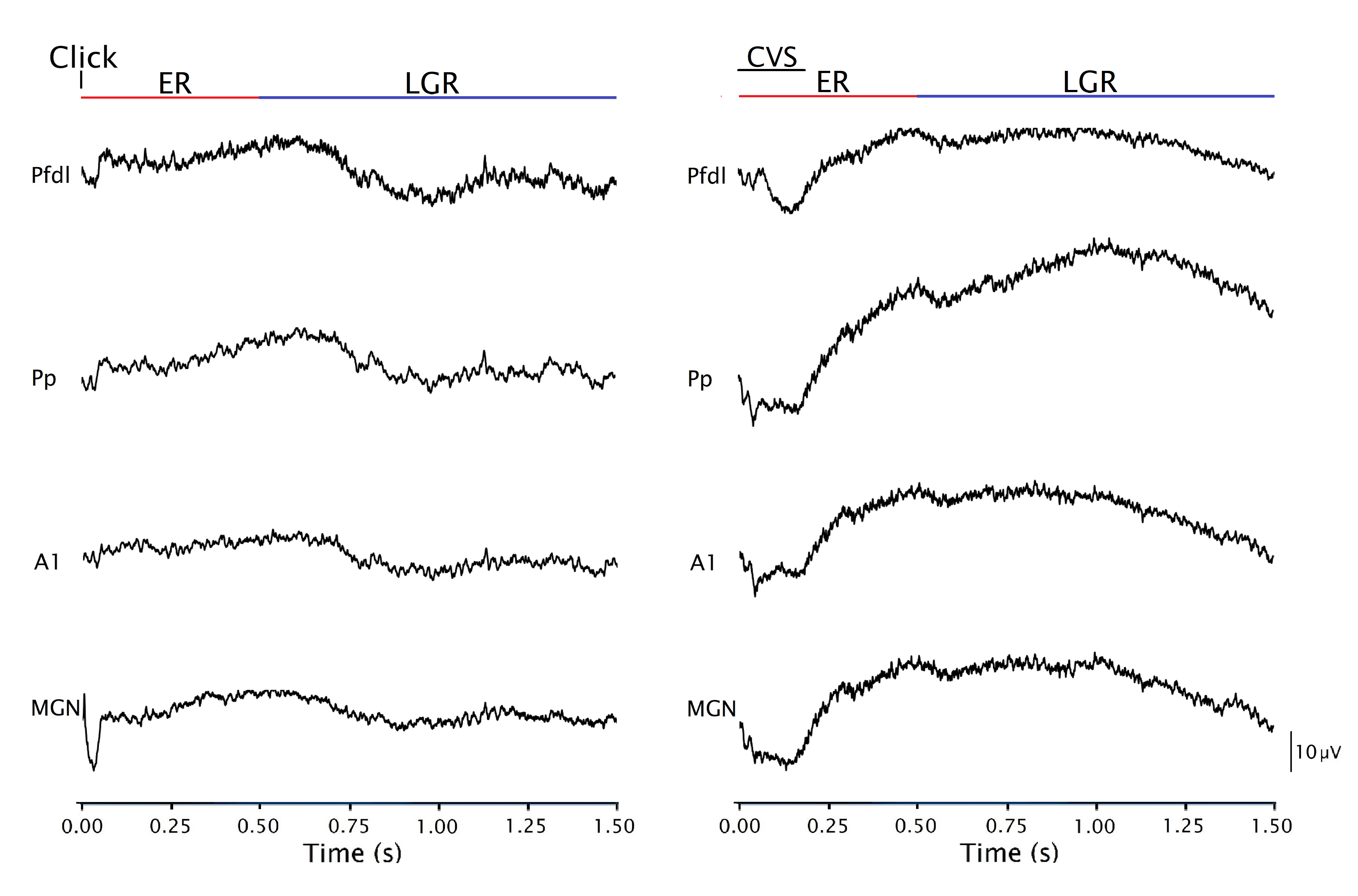

**Supplementary Figure 2.** Averaged recordings of 50 click and CVS stimuli from a representative cat. Evoked potentials in the early response (ER) are clearly observed, but the late gamma response (LGR) is diminished due to averaging. Pfdl: Dorsolateral prefrontal cortex; Pp: Posterior parietal cortex; A1: Primary auditory cortex; MGN: medial geniculate nucleus.

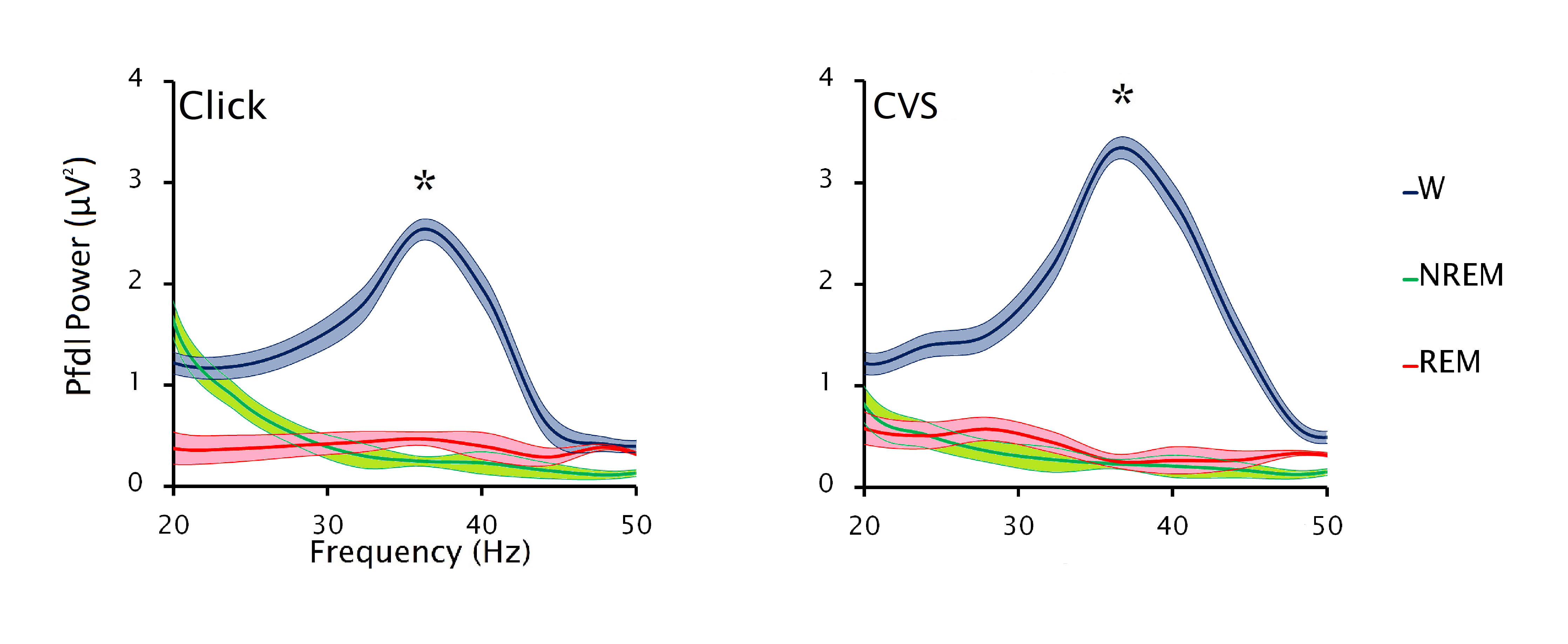

**Supplementary Figure 3.** **Gamma response induced by auditory stimulation during wakefulness and sleep.** Mean and standard error of gamma-band power during the late gamma response (LGR) following click and CVS stimulation during wakefulness, NREM sleep, and REM sleep from the dorsolateral prefrontal cortex (Pfdl)**.** Asterisks (*) show significance (ANOVA and Bonferroni *post hoc* test) with p < 0.05. These analyses were conducted in four cats.

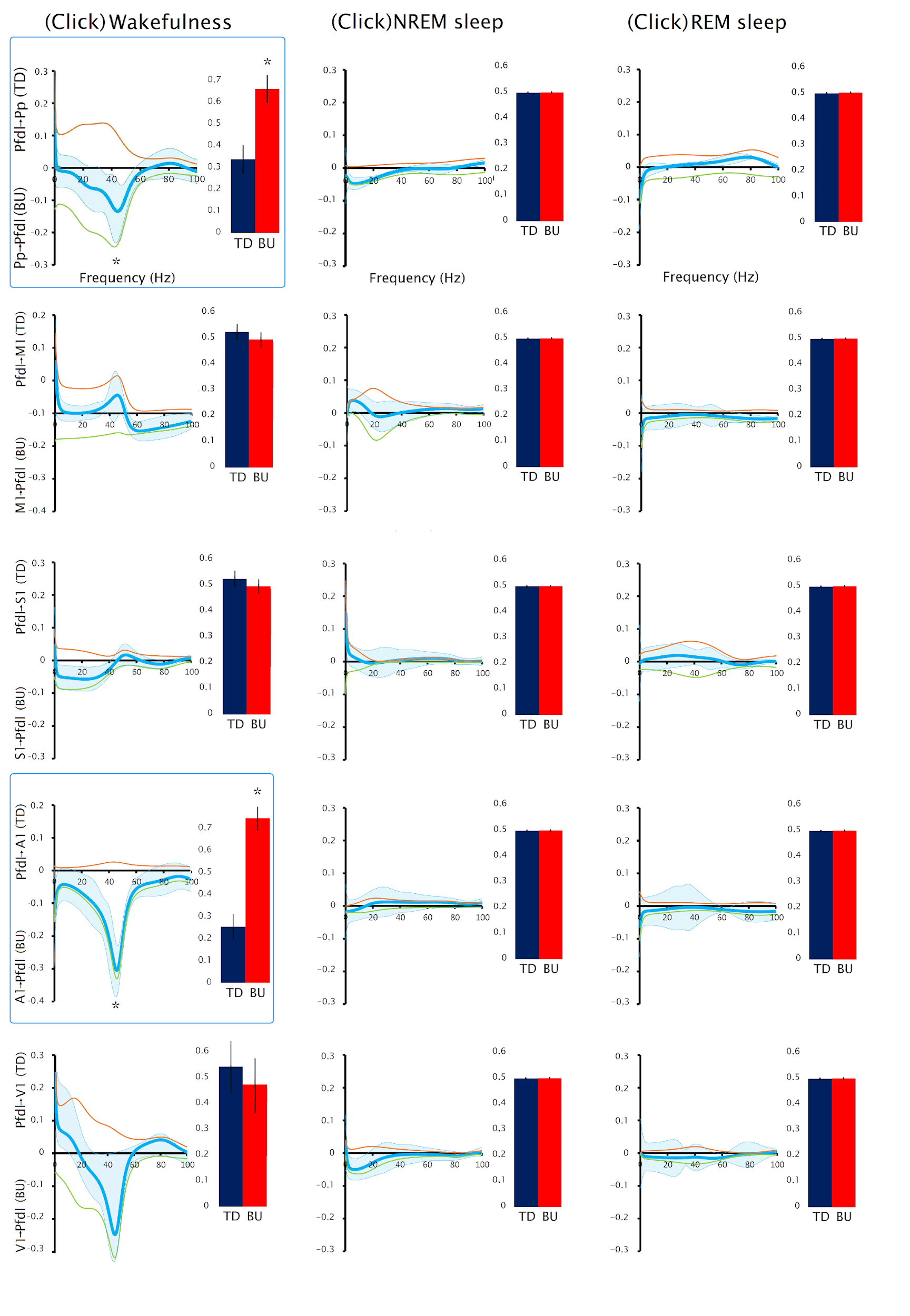

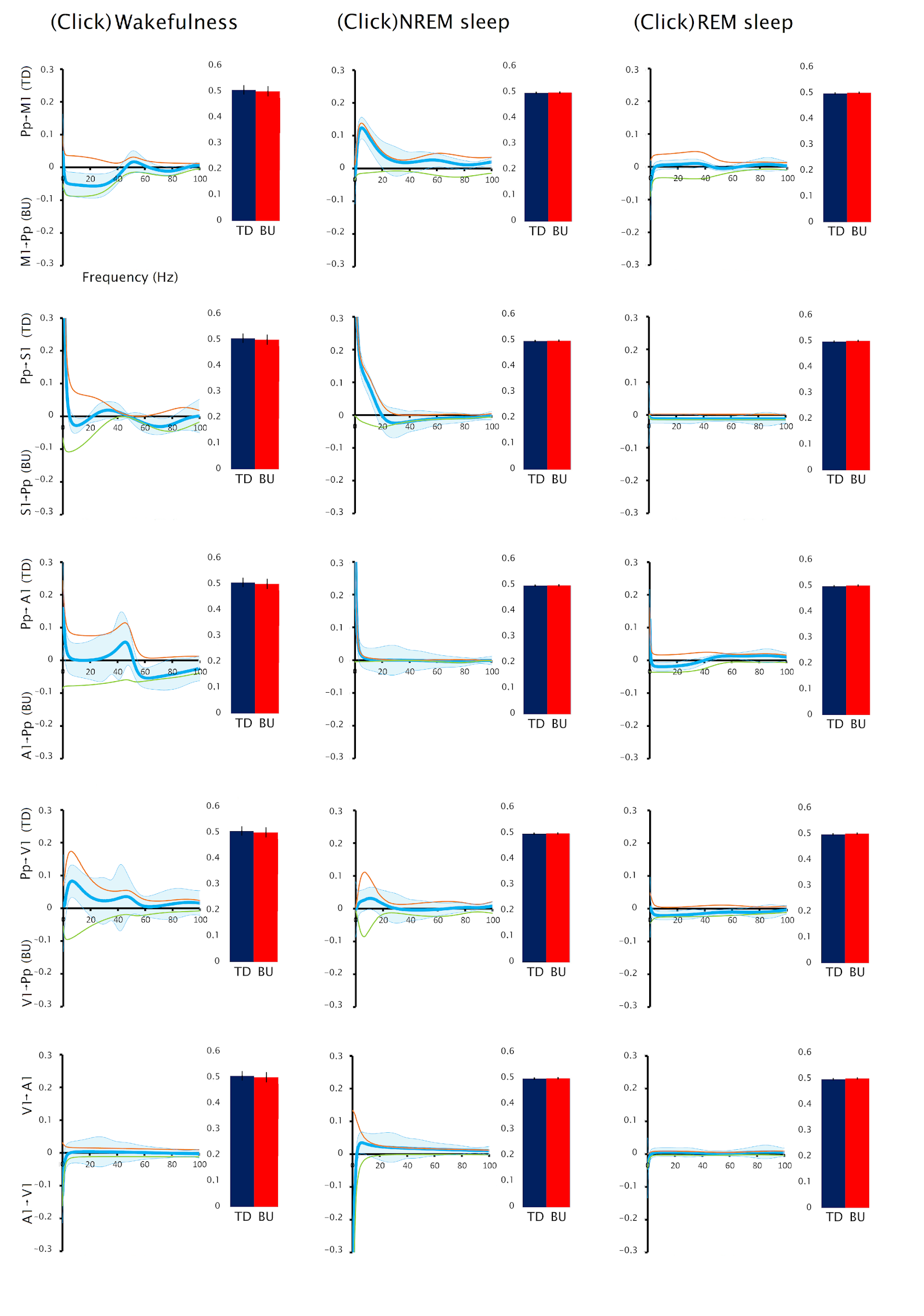

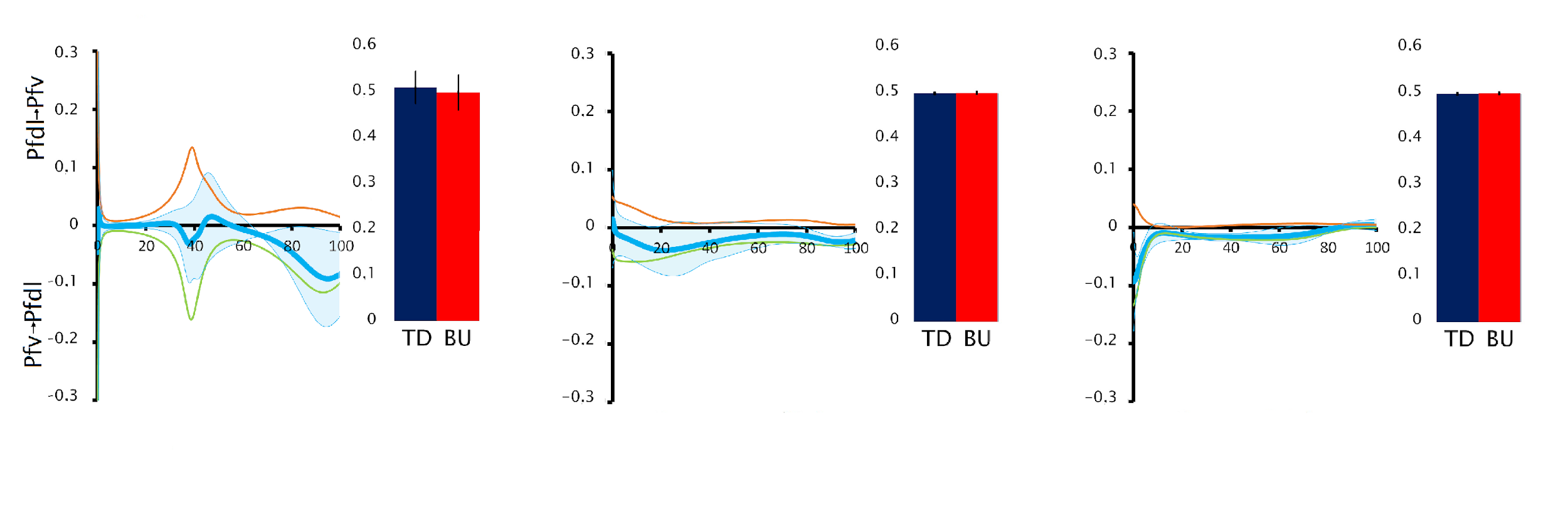

**Supplementary Figure 4.** **Gamma directionality during wakefulness and sleep for all the cortical derivations during LGR induced by click stimuli.** Mean and standard error of Granger Causality spectrum analysis during wakefulness and sleep for all the cortical derivations during LGR induced by click stimuli. The orange line shows the top-down, while the green line the bottom-up directionality. The light blue line illustrates the difference (subtraction) between both directionalities. These analyses are accompanied by graphs that show the number of events of the peaks of the envelopes of the gamma oscillations in each direction. All the analyses were conducted in four cats. Asterisks (*) show significance (ANOVA and Bonferroni post hoc) with all the other time windows (p < 0.05). EEG derivations showing statistically significant differences are outlined by boxes. Pfv: Ventral prefrontal cortex; Pfdl: Dorsolateral prefrontal cortex; M1: Primary motor cortex; S1: Primary somatosensory cortex; Pp: Posterior parietal cortex; A1: Primary auditory cortex; V1: Primary visual cortex. TD: top-down; BU bottom-up

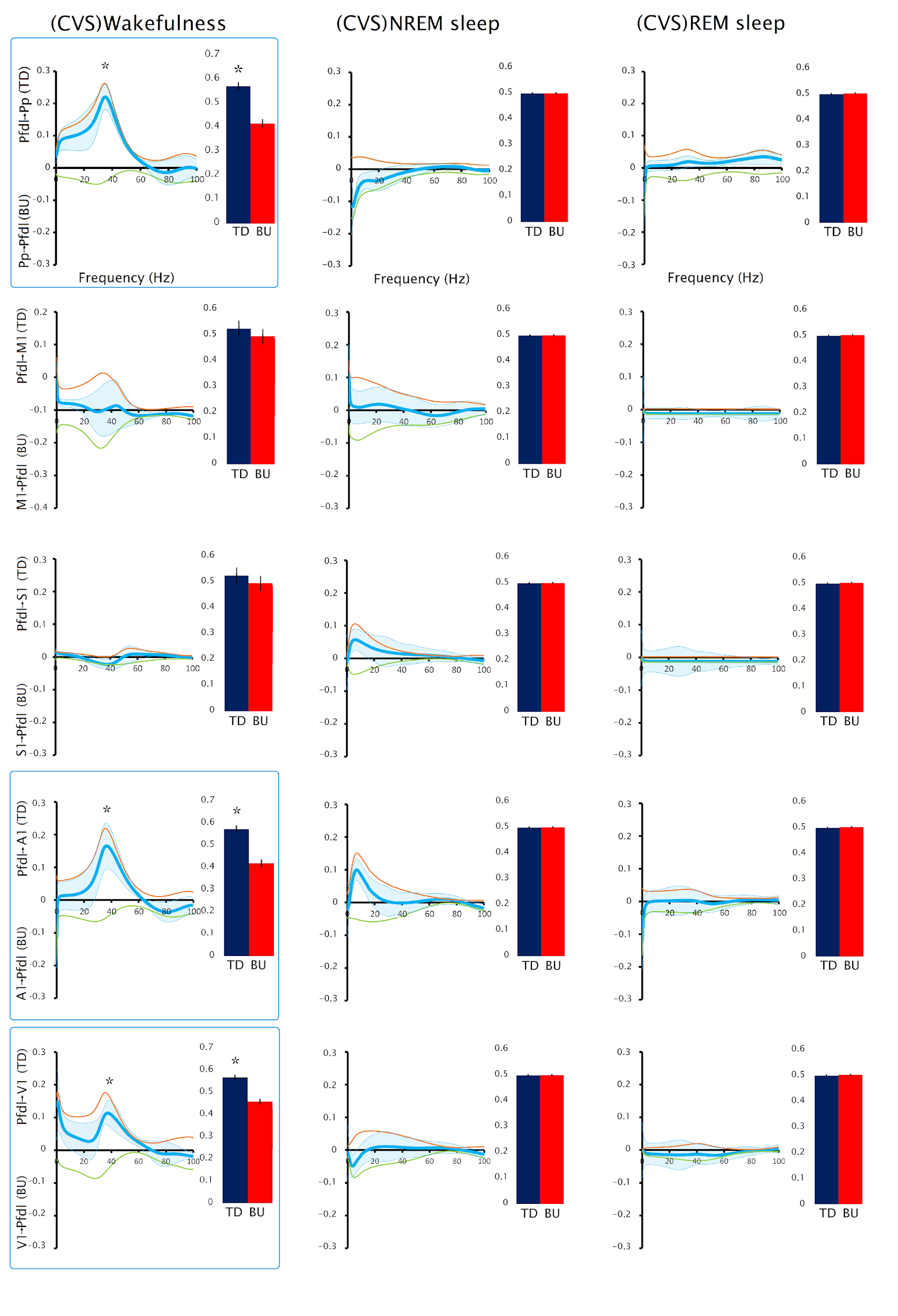

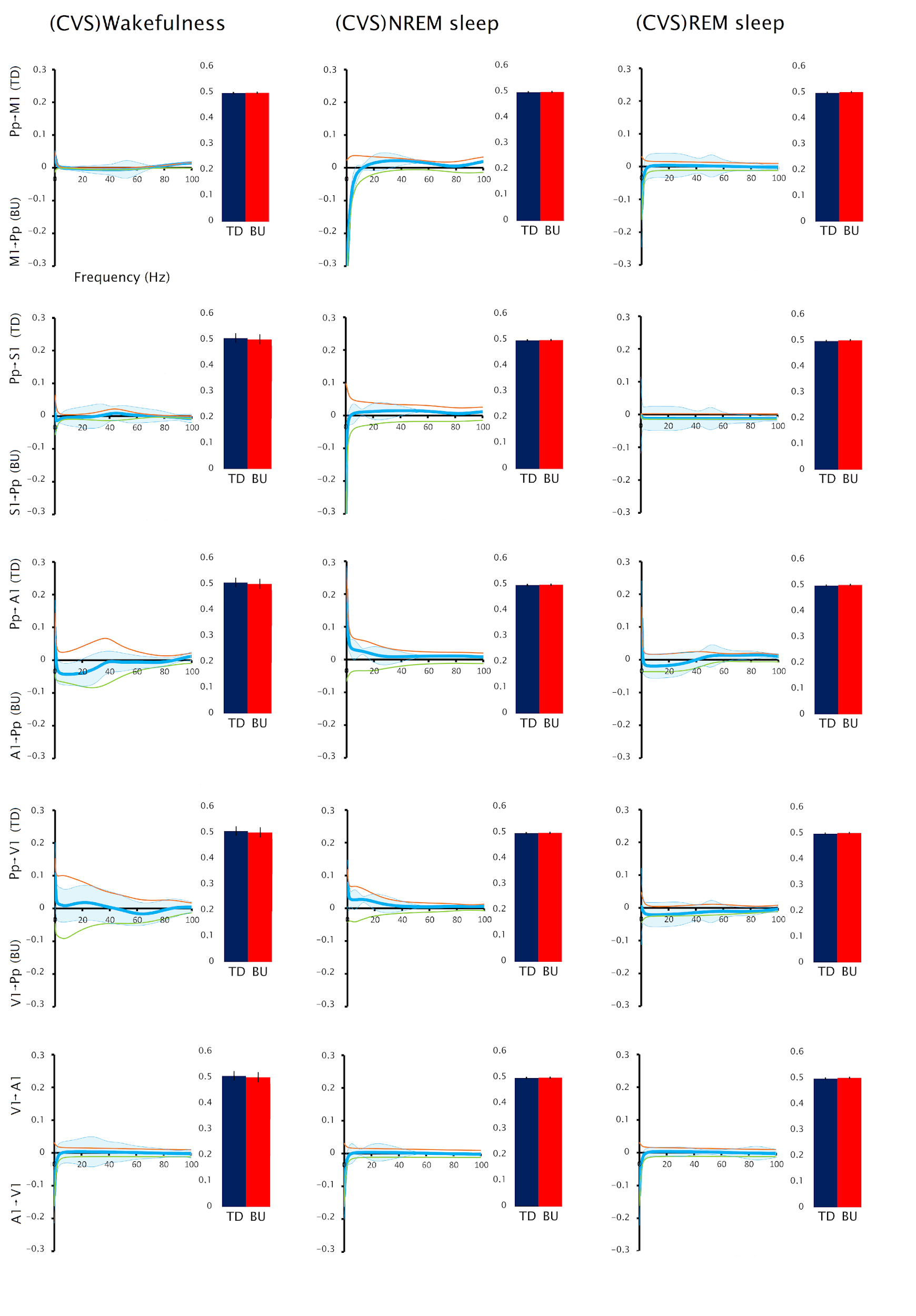

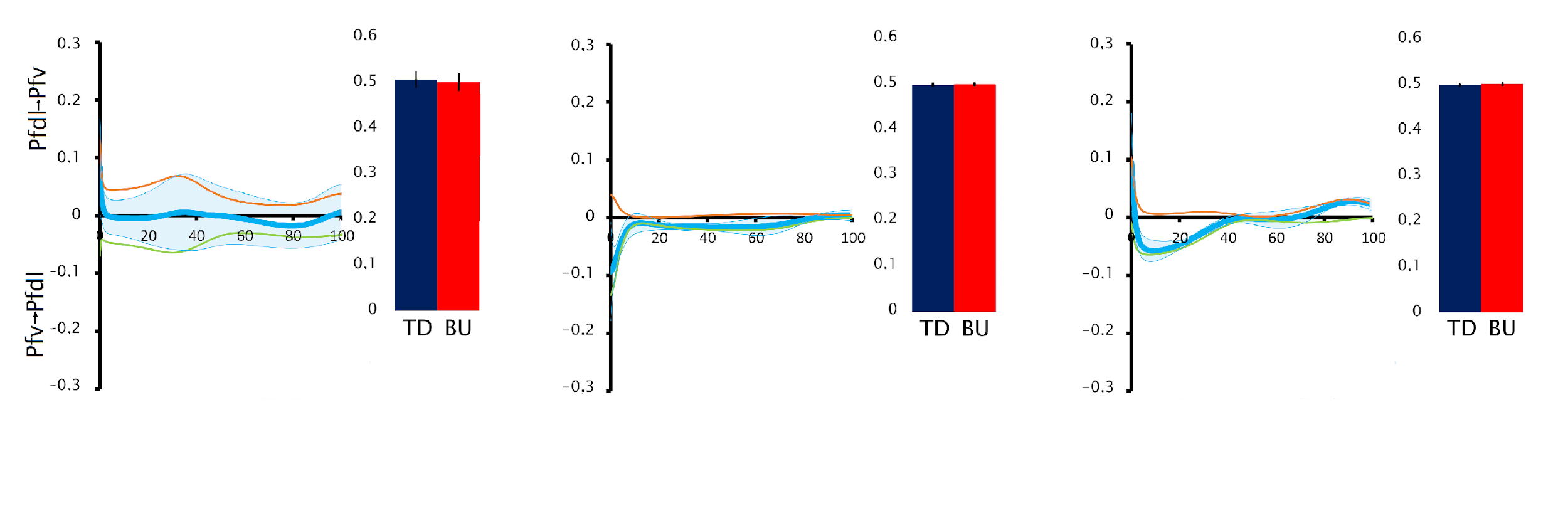

**Supplementary Figure 5.** **Gamma directionality during wakefulness and sleep for all the cortical derivations during LGR induced by CVS stimuli.** Mean and standard error of Granger Causality spectrum analysis during wakefulness and sleep for all the cortical derivations during LGR induced by click stimuli. The orange line shows the top-down, while the green line the bottom-up directionality. The light blue line illustrates the difference (subtraction) between both directionalities. These analyses are accompanied by graphs that show the number of events of the peaks of the envelopes of the gamma oscillations in each direction. All the analyses were conducted in four cats. Asterisks (*) show significance (ANOVA and Bonferroni post hoc) with all the other time windows (p < 0.05). EEG derivations showing statistically significant differences are outlined by boxes. Pfv: Ventral prefrontal cortex; Pfdl: Dorsolateral prefrontal cortex; M1: Primary motor cortex; S1: Primary somatosensory cortex; Pp: Posterior parietal cortex; A1: Primary auditory cortex; V1: Primary visual cortex. TD: top-down; BU bottom-up

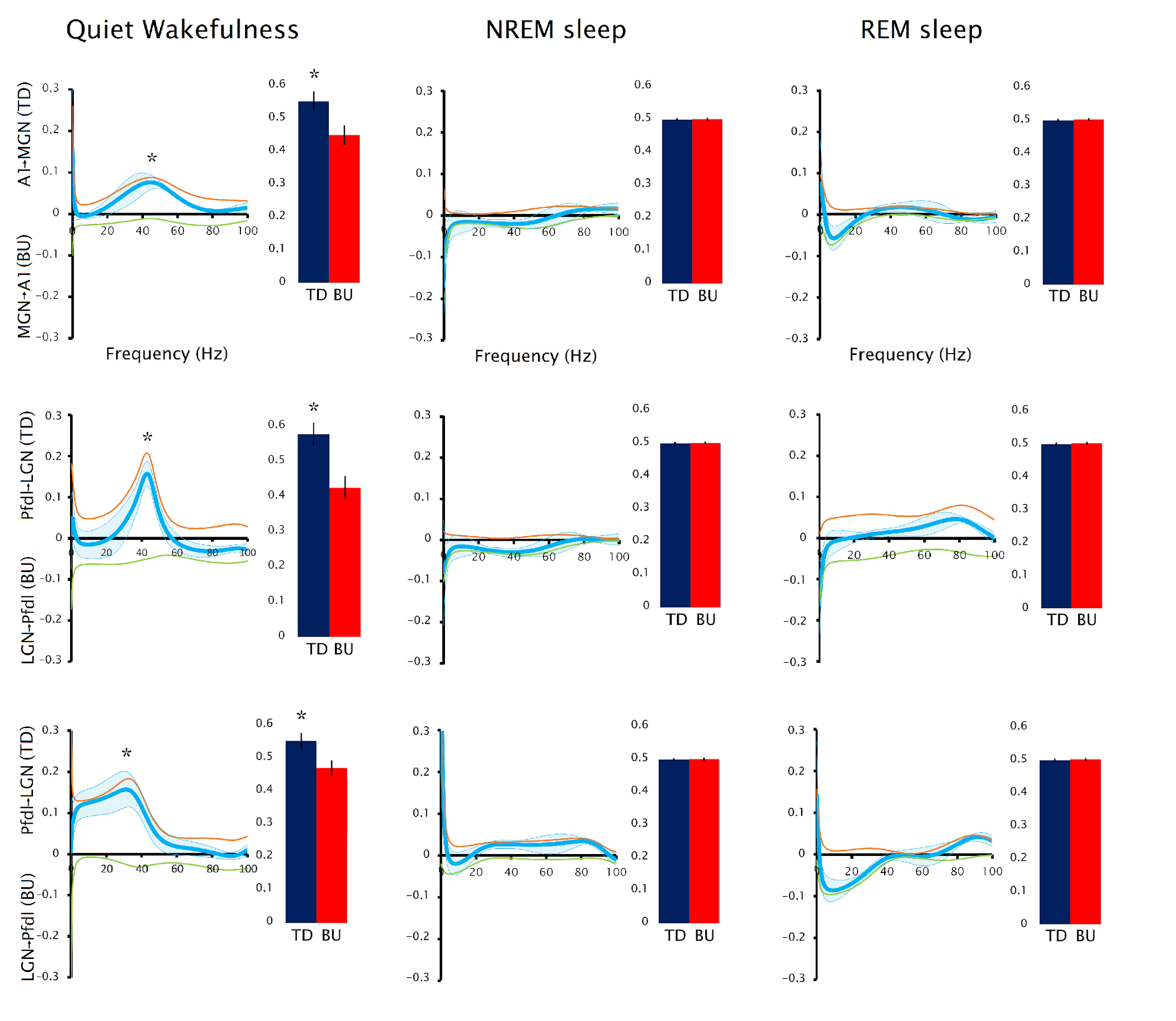

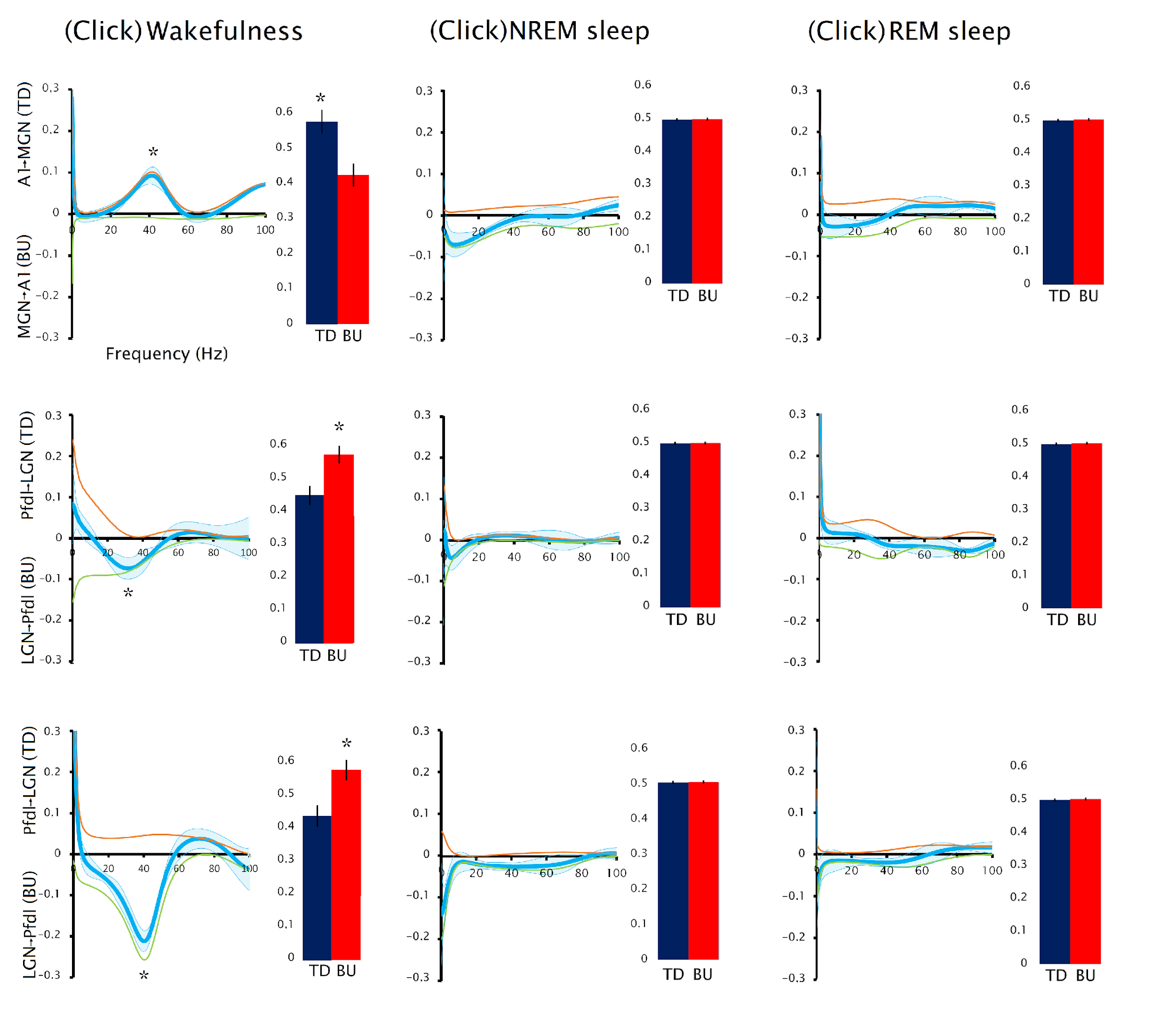

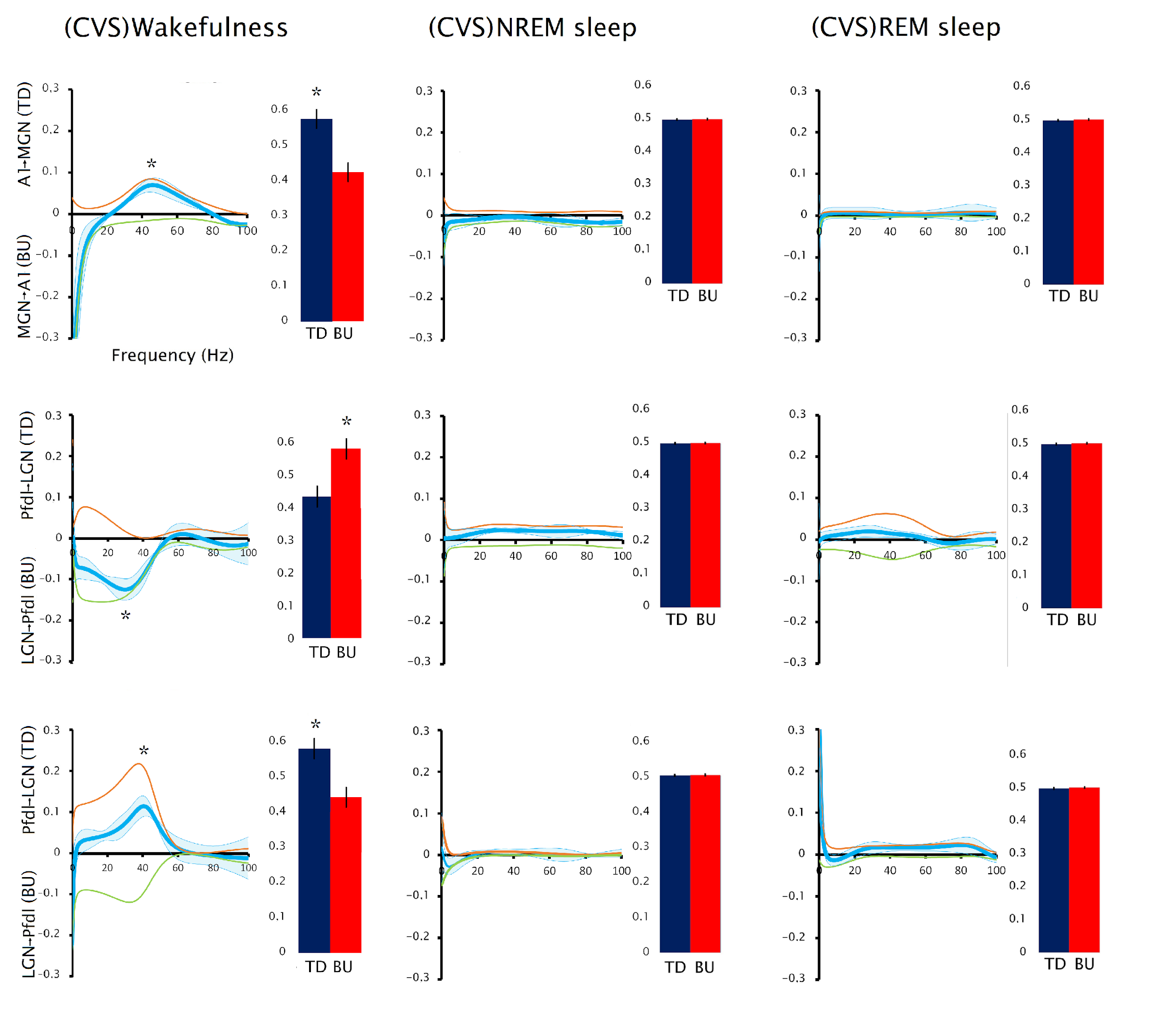

**Supplementary Figure 6.** **Directionality between neocortex and thalamus.** Mean and standard error of Granger Causality spectrum during quiet wakefulness (without sensory stimulation), and during the late gamma response (0.5 to 1.5 seconds after the stimuli) induced by click stimuli or CVS. These analyses are accompanied by graphs that show the number of events of the peaks of the envelopes of the gamma oscillations in each direction.

Asterisks (*) show significance (ANOVA and Bonferroni post hoc) with all the other time windows (p < 0.05). All the analyses were conducted in two cats. TD: top-down; BU bottom-up. Pfdl: Dorsolateral prefrontal cortex; A1: Primary auditory cortex; V1: Primary visual cortex; MGN: medial geniculate nucleus; LGN: lateral geniculate nucleus.

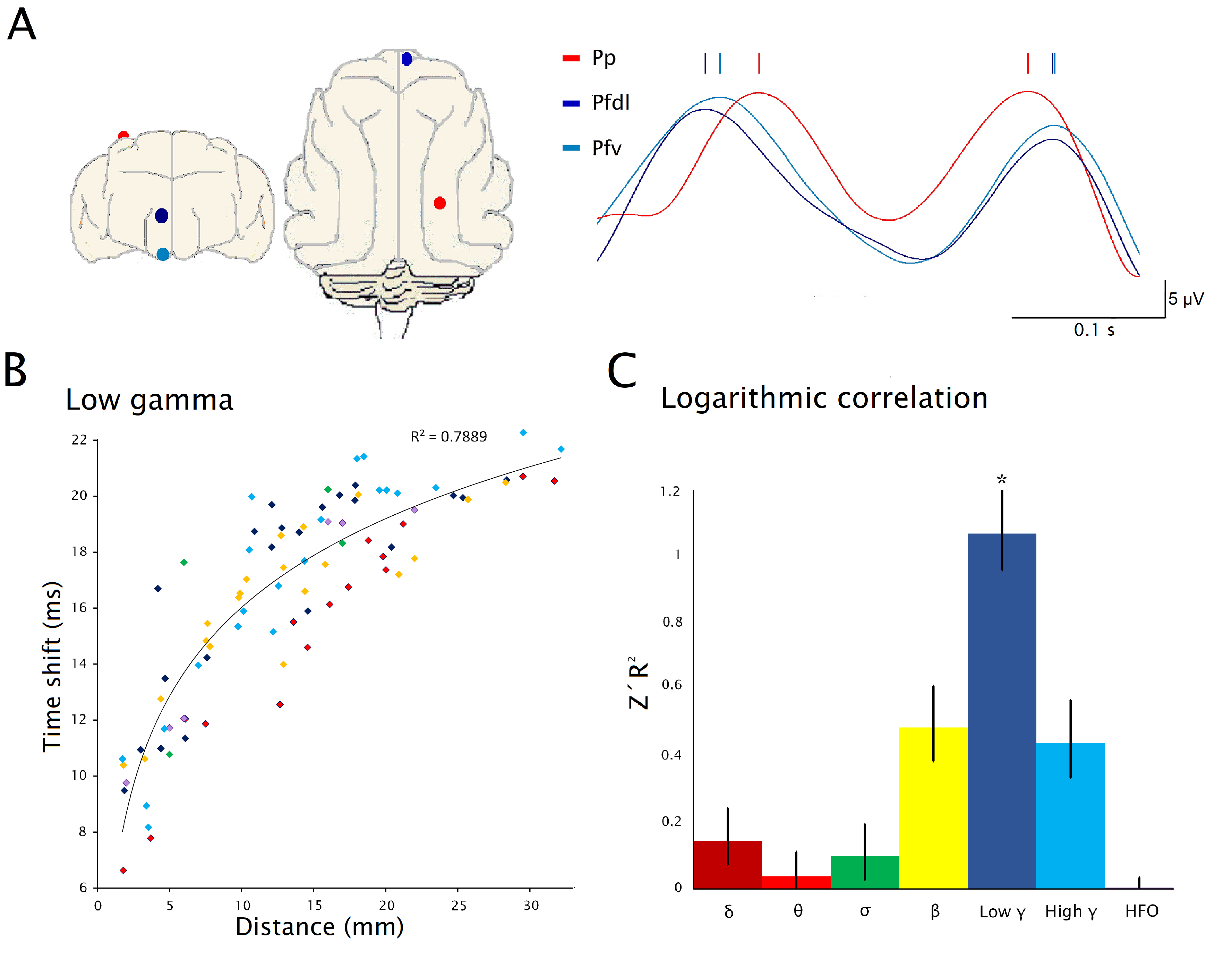

**Supplementary Figure 7.** **Relation between time lag of gamma oscillation and the distance between cortical electrodes during wakefulness.** A. Low gamma oscillations envelopes from dorsolateral prefrontal cortex (Pfdl), ventral prefrontal cortex (Pfv) and posterior parietal cortex (Pp) cortices; the marks indicate the peak of the envelopes. The location of the electrodes is drawn on the left. B. Scatter plot showing the relationship between the mean absolute phase shift of low-gamma envelope peaks and the distance between cortical electrodes. The regression analysis shows that phase shifts correlates with the logarithm of distance between electrodes. C. Fisher's Z-transformed R² values of the logarithmic correlations between envelope-peak phase shifts and the distance between cortical electrodes for the frequency bands analyzed. Correlations were significantly stronger in the low-gamma (low γ) band than in all other frequency bands (p < 0.0001). Data were obtained from six cats.

The precise distance between the cortical electrodes was measured on the cat's skull with a caliper, after the animals were euthanized. The average phase shift for each cortical pair was calculated from the event correlation histograms. Then, we performed a linear and logarithmic regression between these parameters. We compared the regressions obtained of low gamma oscillations (30-45 Hz, low γ) and compare them with the ones obtained from other frequencies such as delta (0.5-4 Hz, δ), theta (4-10 Hz, θ), sigma (10-15 Hz, σ), beta (20-30 Hz, β), high gamma (60-100 Hz, high γ) and high frequency oscillations (100-140 Hz, HFO) using ANOVA and Bonferroni *post hoc* test.
